# Ecological memory preserves phage resistance mechanisms in bacteria

**DOI:** 10.1101/2020.02.23.961862

**Authors:** Antun Skanata, Edo Kussell

## Abstract

Defense mechanisms against pathogens are prevalent in nature, and their maintenance is critical for long-term survival of a species. Such mechanisms, which include CRISPR-mediated immunity in bacteria and the R genes in plants, carry substantial costs to organisms and can be rapidly lost when pathogens are eliminated. How a species preserves its molecular defenses despite their costs, in the face of variable pathogen levels, and across an ecology of localized patches remains a major unsolved problem in epidemiology and evolutionary biology. Using techniques of game theory and non-linear dynamical systems, we show that by maintaining a non-zero failure rate of immunity, hosts sustain sufficient levels of pathogen across an ecology to select against loss of the defense. This *resistance switching strategy* is evolutionarily stable, and provides a powerful evolutionary mechanism that maintains host-pathogen interactions and enables co-evolutionary dynamics in a wide range of systems.

---

Survival of species in the presence of pathogens requires effective defense mechanisms, which exist in a wide range of biological systems [1–4]. Host-pathogen interactions are subject to the availability of susceptible hosts that sustain a viable pool of pathogens, while defense mechanisms are under evolutionary pressure to reduce or eliminate the ability of pathogens to proliferate. Once a threat is removed so is the pressure to preserve the relevant defense mechanism which may carry significant fitness costs [5–7]. In the absence of defenses, pathogens can reemerge and wipe out a host population. The selective forces that drive the defense mechanisms to become highly efficient may thus eventually lead the host to extinction. How species avoid this inherent fragility of defenses against pathogens is not presently understood [8].

Bacteria and their pathogens, the bacteriophages, present a powerful system to study this question. A bacteriophage infects a bacterial cell by attaching to the cell surface, injects its genetic material, replicates and assembles phage particles, and is subsequently released by the cell (Fig. 1). Bacteriophage resistance mechanisms [1] exhibit a great deal of variety across two major classes: (i) *preventative defenses*, which operate by blocking phage receptors [9, 10] or by producing competitive inhibitors [11, 12], and (ii) *immune defenses*, such as restriction-modification systems [13] and CRISPR [14] which cleave phage DNA after it enters the host. In all instances, there is a metabolic cost associated with maintaining the resistance mechanisms [15–17], with the additional fitness costs due to self-targeting in RM systems [18] and CRISPR [19]. Yet, the diversity of phage-host systems in terms of routes of infection and modes of resistance points to strong evolutionary pressures favoring the emergence and maintenance of resistance [2, 20, 21].

**FIG. 1.**
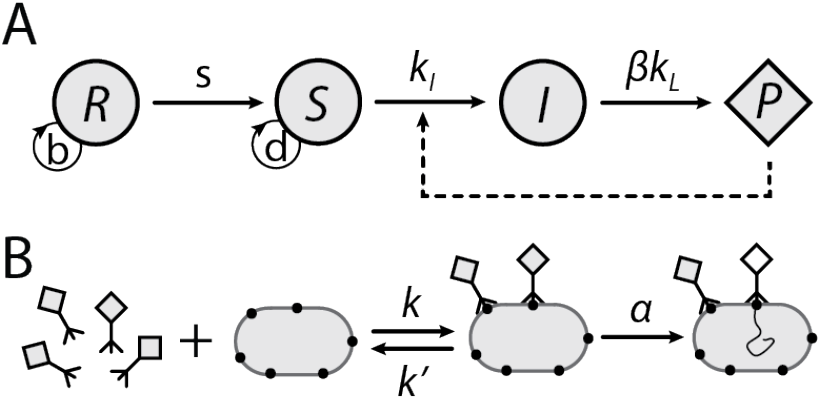
Phage-host population dynamics and molecular interactions. (A) Population dynamics for a preventative defense model. The resistant phenotype *R* generates phage-sensitive cells *S* at rate *s*, on which the phage *P* grows. *I* labels cells infected by phage. Arrows indicate rate constants for each type of transition; the circular arrows indicate the growth rates of *R* and *S* cells, with *d* − *b >* 0 being the cost of resistance. For immune defense, all phenotypes express receptors and absorb phage, but only sensitive cells transition to infection and lysis. (B) Molecular interactions. The phage diffuses onto a host that expresses phage receptors (solid circles), binds reversibly to a receptor and infects the cell by injecting its DNA. Reaction rates are shown on arrows; the phage binding and unbinding rates *k* and *k*′ and phage absorption rate *α* are defined per receptor (see *Appendix*).

A well-studied example of a preventative defense is found in the host-pathogen interaction of *E. coli* and the λ phage, which attaches to the host cell through the LamB transmembrane protein. Preventative resistance is conferred through mutations that cause repression of *lamB*, inducing growth defects on maltose [22] but providing resistance to λ. A subset of these mutations, which involve short amplifications, revert at frequencies of 10^−5^-10^−4^ per division, leading to spontaneous induction of a phage-sensitive phenotype in an otherwise resistant population [23, 24]. In these *resistance switching* strains, phage persist at low frequencies, while in strains that do not switch off their resistance phage are absent. An absence of phage can lead to rapid loss of resistance, for example by derepression of the phage receptor, thereby decreasing costs. Large bacterial population sizes ensure the existence of phage-sensitive mutants, which will sweep to fixation if phage are eliminated.

To study the maintenance of defenses, we model a bacterial host that has already established a phage resistance mechanism and determine the conditions in which resistance is preserved. Remarkably, we demonstrate that spontaneous loss of resistance in single cells – which enables phage to persist in the host’s environment – acts to protect the resistance mechanism from invasion by sensitive mutants. The same strategy allows a host to sustain multiple phage types within a bacteria-phage ecology. We show that coexistence between resistant and sensitive cells strongly depends on the type of defense mechanism, i.e. immune or preventative, and on the resistance switching strategy. Using a combination of ecological dynamics and game-theory models we demonstrate that resistance switching is an evolutionarily stable strategy (ESS) that can be naturally evolved.

## I. RESULTS

To study the evolutionary stability of resistance mechanisms, we first determine the conditions in which resistant hosts coexist with pathogens, and then consider bacterial invasions that take place in an ecological setting. We model the interactions between phage (*P*) and bacteria that express a resistant phenotype (*R*) or a sensitive phenotype (*S*). Phage infect sensitive cells, generating infected cells (*I*) at rate *k*_*I*_ , and these lyse at rate *k*_*L*_ to produce *β* new phage particles; *βk*_*L*_ is the phage *burst rate*, i.e. the overall rate at which phage particles are released into the environment. We model the latency period by explicitly including the infected phenotype in our equations [25, 26]. Resistance switching is modeled as spontaneous conversion of *R* cells into *S* cells at rate *s*, while the cost of resistance is given by the difference between resistant and sensitive cells’ growth rates. Figure 1A shows these population dynamics schematically and the equations are given below, where capital letters denote the population size of each component of the system:

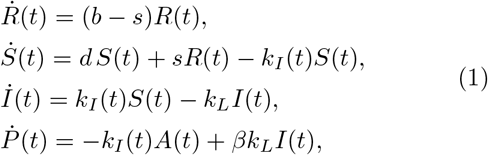

where *k*_*I*_ (*t*) is the host infection rate (see *Appendix* and *Supplementary Materials*), *d* and *b* are growth rates of *S* and *R*, and *A* is the total number of phage-absorbing hosts. For preventative defenses, only the sensitive and infected phenotypes absorb phage, and we set *A* = *S* + *I*. For immune defense systems, all phenotypes absorb phage, and *A* = *R* + *S* + *I*.

The nature of the phage-bacteria interaction, which is mediated at the molecular level by phage receptor kinetics and other processes, determines the dynamics at the population level. It is therefore critical to consider how molecular details are incorporated into the cellular population dynamics of Fig. 1A. Figure 1B shows the coarsegrained mechanism of infection by a phage freely diffusing onto a host cell [27–29]. A phage-receptor complex is formed reversibly, followed by irreversible injection of phage genetic material into the cell at rate *α*, which we refer to as the phage *absorption rate*. The overall infection rate *k*_*I*_ per host cell is a function of *P* and *A*, and depends on the average number of phage receptors per cell, *n*_*r*_, and on the molecular rate constants shown in Fig. 1B (see *Appendix* and further details in *Supplementary Materials*). Phage burst sizes vary by orders of magnitude among different phages grown on similar hosts (e.g. *β* ≃ 10^2^ − 10^4^ in *E.coli* [30]) and a tradeoff in burst rate of the form *βk*_*L*_ ≈ constant [31, 32] indicates that changes in *k*_*L*_ may be compensated by changes in *β*.

To fully specify the population model requires a relation between the external environment and the growth of host cells, which can be incorporated in Eq. (1) in different ways (*Supplementary Materials*). This organism-environment interaction can take many different forms, and its key role is to set bounds on the total population size in models that otherwise admit exponentially growing solutions. We find that the specific growth modality does not impact the major outcomes below. We present results for growth in rich media with feedback dilution (turbidostat growth) in the main text, or in a nutrient-limited environment (chemostat growth) in *Supplementary Materials*.

To characterize the population dynamics of the phage-bacteria system, we solve the model equations and determine the long-term population structure. Regardless of the initial numbers of phage and bacteria, the population as a whole approaches a set of stable outcomes, or *phases*, in which the population composition either remains fixed or exhibits periodic oscillations. We show below how the particular phase depends on the phage’s burst and absorption rates, and identify the range of phages for which resistance is stable within a single population. To determine the conditions under which a population preserves resistance to phage, we examine the invasion dynamics between different bacterial strains, e.g. resistance switching strains with different switching rates (*s* > 0), non-switching resistant strains (*s* = 0), or sensitive strains. Using a layered modeling approach, spanning from molecular to population to ecological levels, we determine the stable evolutionary outcomes for a minimal ecology consisting of a large number of independent populations.

### A. Stable phases of phage-host systems

The state of the phage-bacteria system can be visualized as a ternary plot in which each corner corresponds to a monomorphic population composed entirely of sensitive cells, resistant cells, or phage particles (Fig. 2A). Flow lines within the diagram show the solutions of the model starting from different initial conditions. Points in the interior of the triangle represent different number compositions of host cell phenotypes and phage, using total counts of cells and phage for normalization; one can alternatively plot the composition as biomass fractions, which is a one-to-one transformation of the ternary plot that preserves all topological features including fixed points, trajectory structure, and stability (*Supplementary Materials*). We present results separately for the preventative and immune defense models below.

**FIG. 2.**
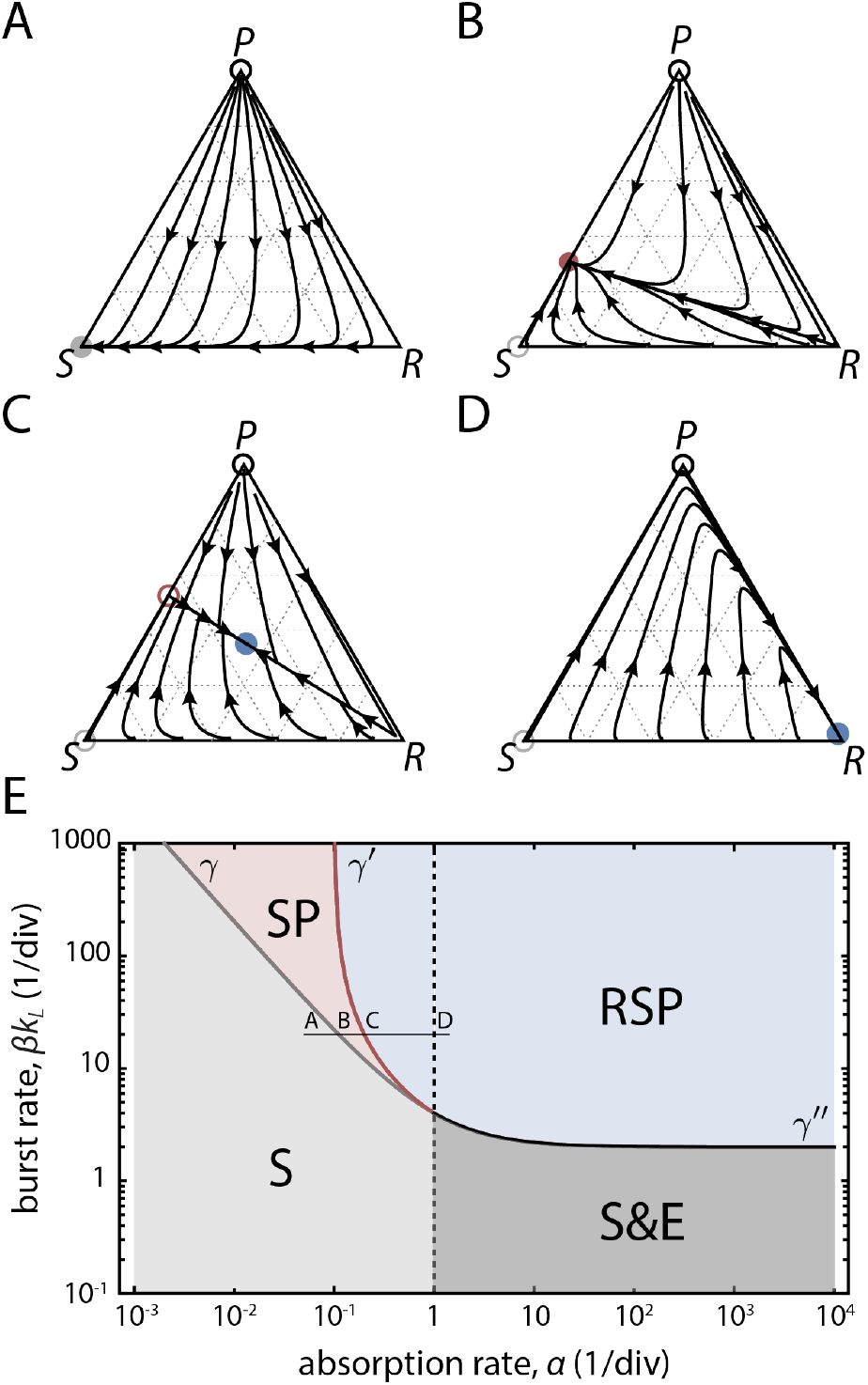
Stable phases for preventative defenses. (A)-(D) Flow diagrams of Eq. (1), plotting frequencies of the resistant (*R*) phenotype along the bottom edge, sensitive (*S*) phenotype along the left edge and phage and infected (as *P* + *I*) along the right edge in a ternary plot. Empty (filled) circles - unstable (stable) fixed points. Color of the fixed point matches the phase displayed in the phase diagram below. Host extinction at the top corner is represented with a black empty circle. (E) The phase diagram of preventative defenses as a function of burst rate *βk*_*L*_ and phage absorption rate *α*. The S phase (gray region) carries only the *S* phenotype, the SP phase (pink region) carries the *S* phenotype and phage, the RSP phase (blue region) carries all phenotypes and phage, the S&E phase (dark gray region) corresponds to bistability between phase S or host extinction E. The curves *γ* (gray), *γ*′ (dark red) and *γ*″ (black) denote locations of bifurcations. Letters A-D represent locations for which we plotted flow diagrams in the corresponding panels above. All rates are in units of 1/division, with *d* = 1, *b* = 0.9, *k*_*L*_ = 1 and *s* = 10^−4^. Results are shown for high phage-receptor binding affinity (*κ* → 0) and minimal sensitivity (*n*_*r*_ = 1). See *Supplementary Materials* for *κ >* 0 and Figs. S6 – S8 for dependence on *k*_*L*_, *n*_*r*_ and *b*.

#### 1. Preventative defenses

Representative flow diagrams for the preventative defense model, in which resistant cells do not absorb the phage, are shown in Fig. 2A-D, where each panel corresponds to a different phage absorption rate *α* and burst rate *βk*_*L*_. Across all possible combinations of phage parameters, there are four possible long-term outcomes of the dynamics which correspond to fixed points of the model equations: (i) fixation of *S* [S phase], (ii) coexistence of *S* and *P* [SP phase], (iii) coexistence of *R*, *S*, and *P* [RSP phase] and (iv) host extinction [E phase].

Depending on the phage parameters, one or more of these phases may be stable to small perturbations of the population composition. The stable phases are shown as distinct regions in the space of phage parameters (Fig. 2E), separated by curves *γ* and *γ*′, which correspond to transcritical bifurcations of the dynamical system, and a curve *γ*″ where the RSP fixed point becomes unstable (see *Supplementary Materials* and Figs. S3 & S4). There also exist regions where two phases are stable (for *α* > *d*/*n*_*r*_), and which phase is observed depends on initial conditions; these include the S&E bistable region and a narrow region of RSP&S bistability (*Supplementary Materials*, Fig. S4). The phase diagram for a model that lacks the resistance phenotype is shown in *Supplementary Materials*, Fig. S1.

In the S phase, which is the unique stable phase for *α* < *d*/*n*_*r*_ and *βk_L_ < γ*, the resistant phenotype and phage will be outgrown by the sensitive phenotype (Fig. 2A). As *α* increases along the thin black horizontal line in Fig. 2E, it crosses the curve *γ*, where the phage can coexist with the host in the SP phase (Fig. 2B). In this phase, the growth rate of *S*, while reduced by the phage, is still larger than the growth rate of *R*, and resistance cannot establish. With further increase in *α*, the growth rate of *S* cells decreases until it equals the growth rate of *R* at the location of *γ*′, where the system transitions to the RSP phase (Fig. 2C). Beyond *γ*′, the frequency of *R* increases while the frequencies of *S*, *I* and *P* decrease to a stable fixed point located in the interior of the simplex, near the *R* corner in Fig. 2D. Since in the RSP phase the dynamics between *S*, *I* and *P* are generated by phenotype switching from *R*, the growth rate of the entire host-phage system is equal to that of the *R* pheno-type, or *b − s* (*Supplementary Materials*). Interestingly, in this phase a phage that infects at a higher rate will be present at lower frequency, because resistance will have a higher selective advantage in the presence of a stronger pathogen (*Supplementary Materials*, Fig. S3).

In the absence of switching (*s* = 0), the RSP phase reduces to a pure resistant population. Any amount of non-zero switching (*s* > 0) will generate *S* on which phage can grow, and promote phage presence in the host population. Furthermore, in *Supplementary Materials*, Fig. S5 we show that non-zero switching allows distinct phage strains with different parameters to coexist within the same host population. We determine when a strategy with *s* > 0 is evolutionarily stable below (see Sec. I C).

#### 2. Immune defenses

Representative flows and the phase diagram for immune defenses, in which all phenotypes absorb the phage, are shown in Fig. 3A-C. Absorption of phage by the resistant cells directly couples phage and resistant subpopulations in equations (1) and generates a region where none of the four possible fixed points are stable. In this region, stable limit cycles are possible in which *R*, *S*, and *P* levels oscillate periodically (Fig. 3C, white region); this region is therefore part of the RSP phase. In Fig. 3A and 3B we show two such orbits, one for the dynamics located near the edge of that region and one located further inside the region. The limiting orbit in Fig. 3B passes extremely close to the *P* = 0 boundary, a behavior that in finite systems would eventually lead to loss of phage and collapse to a stable S fixed point. By increasing the switching rate, such orbits are pulled away from the boundary and toward the interior of the simplex (*Supplementary Materials*, Fig. S10).

**FIG. 3.**
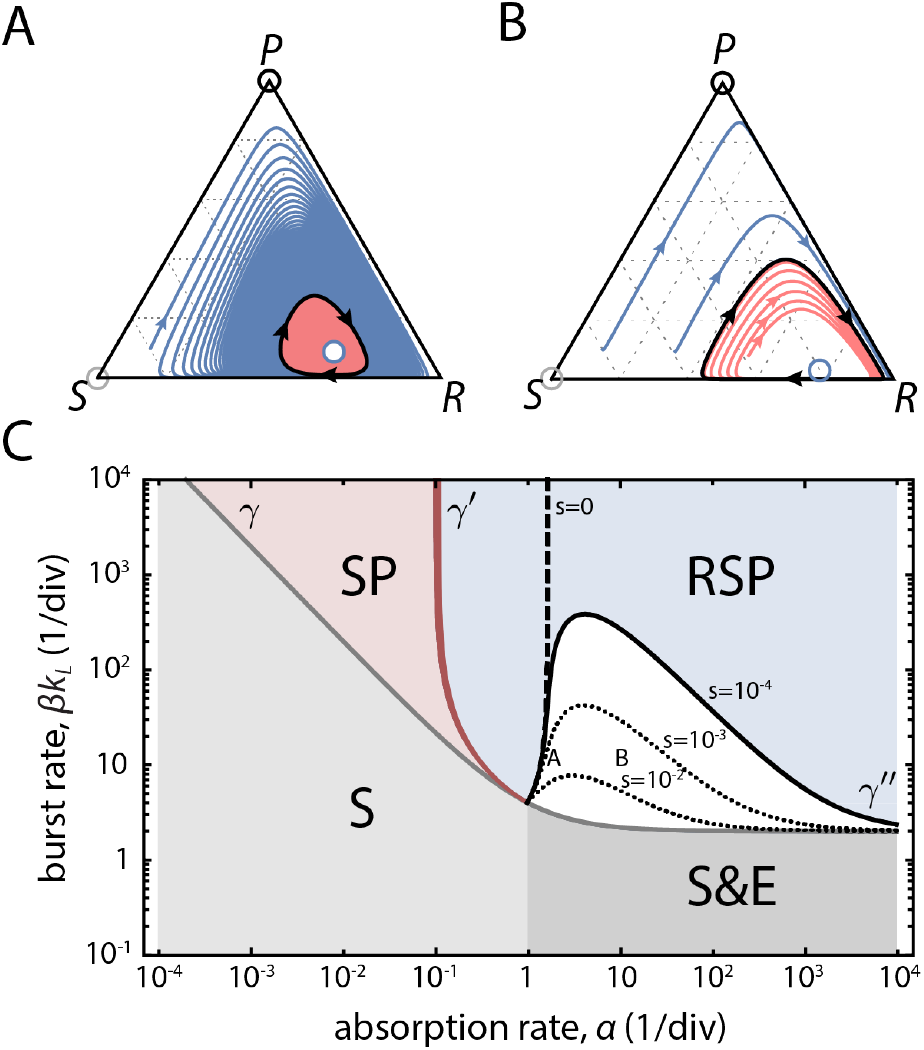
Stable phases for immune defenses. (A)-(B) Flow diagrams of Eq. (1) applied to immune defenses show periodic orbits occurring in the RSP region of the phase diagram shown in panel C. Blue (pink) curves approach the orbits from two directions. The blue empty circle marks the location of the unstable interior fixed point. (C) The phase diagram of immune defenses as a function of burst rate *βk*_*L*_ and phage absorption rate *α*. The S, SP, RSP and S&E phases are defined as in Fig. 2. Curves *γ* (gray), *γ*′ (dark red) and *γ*″ (black) denote locations of bifurcations. The white region bounded by *γ*″ drawn for *s* = 10^−4^ contains no stable fixed points, and admits a stable limit cycle. Dotted black curves denote the *γ*″ curve for higher switching rates; the dashed black curve corresponds to *γ*″ for *s* = 0. Letters A and B denote approximate locations on the phase diagram for which we plotted flow diagrams in the corresponding panels, for *s* = 10^−3^. Remaining parameters are as in Fig. 2.

A Hopf bifurcation curve *γ*″ separates the stable and unstable fixed points of the RSP phase, and is shown in Fig. 3C for different values of *s* > 0 (solid and dotted black curves) or for *s* = 0 (dashed black curve). As *s* increases from zero, *γ*″ confines the region of periodic dynamics to lower phage burst rates, while the *γ*′ curve shifts (slightly) to higher *βk*_*L*_ across the thickness of the dark red curve in Fig. 3C.

#### 3. Parameter dependence of phase diagrams

We examined the structure of phase diagrams when model parameters are varied, including *k*_*L*_, *n*_*R*_, *d*, and *b* (*Supplementary Materials*, Figs. S6 – S8). Figures 2 and 3 show results for a minimally sensitive phenotype (*n*_*r*_ = 1), while increasing the number of receptors per cell moves the phase boundary *γ*′ separating SP and RSP phases to the left, expanding the domain of the RSP phase (*Supplementary Materials*, Fig. S7). Decreasing the cost of a defense mechanism given by the growth rate difference *d − b* likewise moves the *γ*′ boundary to the left (*Supplementary Materials*, Fig. S8). Both dependencies can be seen from the exact expression for *γ*′, which has a vertical asymptote at *α* = (*d*−*b*+*s*)/*n*_*r*_ (*Appendix*). The phase boundary *γ* separating RSP from S and E phases is independent of *n*_*R*_ and *b* (*Appendix*). We additionally examined the possibility that a phage that injected DNA into a cell remains bound to the receptor and blocks it to further phage absorption (*Supplementary Materials*, Fig. S9). Phase diagrams were qualitatively unchanged when the phage lysis rate was varied (*Supplementary Materials*, Fig. S6), since *k*_*L*_ affects only the overall scale in the expressions for *γ* and *γ*′ (*Appendix*). Likewise, changing the growth rate *d* does not impact the phase diagram, as all rates are expressed in units of cell divisions. Phage are known to decay at rates that are several orders of magnitude lower than cellular division rates [33] and therefore phage decay does not impact the phase structure (*Supplementary Materials*). To assess the importance of density dependent growth on the results, we implemented the model using chemostat growth with a limiting resource, which displays a similar phase structure (*Supplementary Materials*, Fig. S2).

### B. Invasion dynamics in a single patch

We now examine invasions in a single patch or territory occupied by distinct host strain genotypes including a resistance switching strain **R**_**s**_ (phenotype *R* switches to *S* at rate *s* > 0), a non-switching resistant strain **R**_**0**_ (phenotype *R*), and a sensitive strain **S** (phenotype *S*). We consider a single phage type with parameters in the RSP phase, hence only the **R**_**s**_ strain carries the phage, while the **R**_**0**_ and **S** strains do not. We analyze dynamics within a patch dominated by one strain when a second strain is introduced initially at low frequency.

For preventative defenses, Fig. 4A shows that **S** is replaced by **R**_**s**_, as **R**_**s**_ brings phage-infected cells into the patch that lyse to release phage, which then infect the **S** strain and drive it to extinction. However, switching to the sensitive phenotype reduces the growth rate of **R**_**s**_ in the presence of phage, which allows an **R**_**0**_ strain to invade over a timescale 1/*s*, and eventually drive both the **R**_**s**_ strain and phage to extinction. Subsequently, an **S** strain could invade the patch, replacing **R**_**0**_ in the absence of phage. It is therefore crucial to consider how invasion trajectories such as **S** → **R**_*s*_ → **R**_0_ → **S** …, may impact the preservation of resistance at the level of inter-patch dynamics, which we analyze below.

**FIG. 4.**
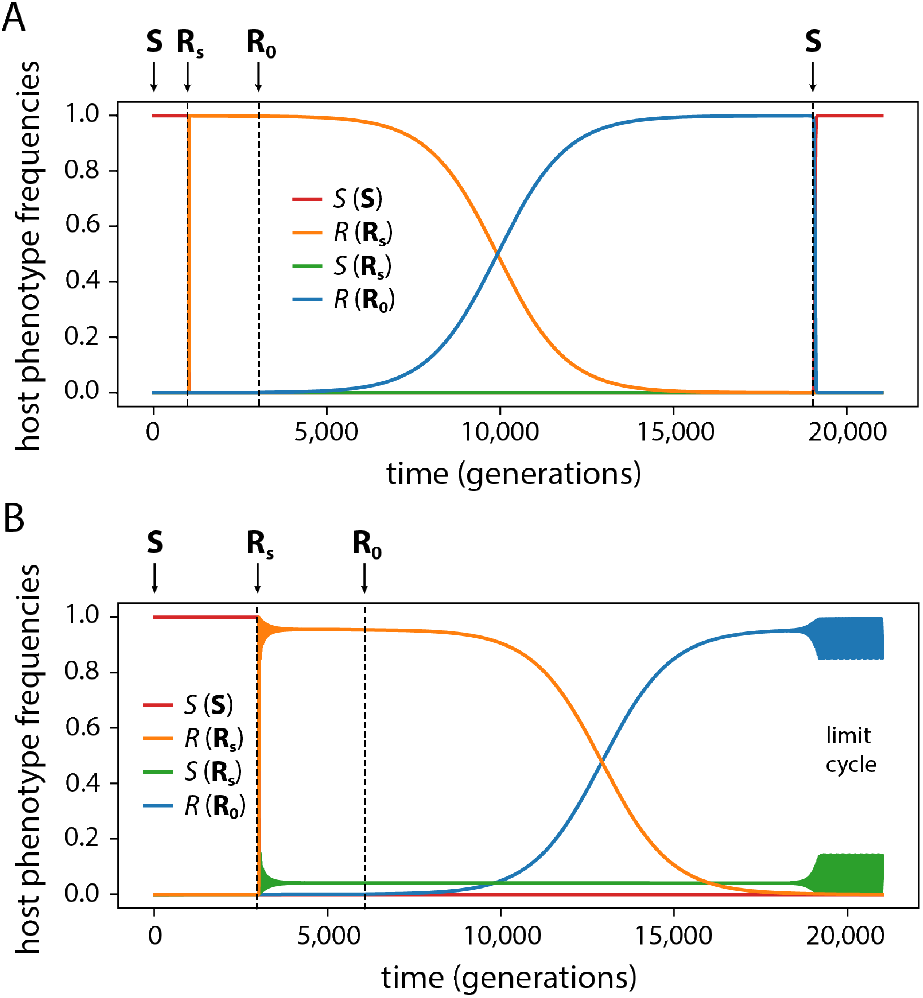
Invasion dynamics for (A) preventative and (B) immune defense systems showing phenotype frequencies in the host population. Arrows indicate times at which an invading strain is added to the population. We label by *R*(**R_s_**) and *R*(**R_0_**) the resistant phenotype of each strain separately, and similarly by *S*(**R_s_**) and *S*(**S**) the sensitive phenotypes. The resistance switching strain **R_s_** outcompetes the sensitive strain **S** as it carries phage which lyse sensitive cells. In (A), **R_0_** invades and eventually replaces **R_s_**, and can then be invaded by **S**. In (B), **R_0_** invades **R_s_**, replacing the switching *R* phenotype while coexisting with the *S* phenotype of the **R_s_** strain and its phage. The RSP fixed point, which was stable for the **R_s_** strain, is unstable for the **R_0_** strain and the dynamics transitions to a stable limit cycle. The model uses turbidostat control with *s* = 10^−3^, *α* = 10, *β* = 50; all remaining parameters are as in Fig. 2.

For immune defenses, the **S** → **R**_*s*_ → **R**_0_ invasion proceeds in a similar way, but the critical difference is that competitions between **R**_0_ and **R**_*s*_ resolve in a surprising manner: the non-switching *R* phenotype of the **R**_0_ strain drives the switching *R* phenotype of the **R**_*s*_ strain to extinction at rate *s*, but the patch reaches a coexistence of **R**_0_, *S* and *P*, either as a fixed point or limit cycle. The coexistence persists despite the fact that there is no new generation of *S*. Instead, **R**_0_ cells act as a phage sink and alleviate the phage pressure on *S* so that its growth rate matches that of **R**_0_, enabling true coexistence.

The stable coexistence of sensitive and immune resistant cells in the presence of phage suggests the possibility that unrelated ‘cheater’ strains could enjoy the benefit of coexistence with immune strains without paying the cost of resistance. Analysis of the stable phases in that scenario (*Supplementary Materials*, Fig. S11) indicates that resistance switching can prevent the establishment of such immune defense cheaters. For higher values of *s*, as can be achieved in the mechanism of CRISPR spacer loss [34], higher levels of phage are present, and an **R**_**s**_ strain generates selection pressure against invaders that is proportional to *βs*. Invaders whose growth advantage is below that threshold will be driven to extinction (*Supplementary Materials*).

### C. Evolutionary stability in an ecology of patches

We now discuss the large-scale dynamics of migration and invasion that take place in the setting of a large number of patches. We introduce the following patch invasion dynamics for a preventative defense system. We initialize the ecology such that each patch is occupied by a single strain, with all strains sufficiently represented, and where phage is always present together with the **R**_*s*_ strain. We index the three possible patch types by *i* ∈ {**R**_*s*_, **R**_0_, **S**}, and let *x*_*i*_ denote the patch frequencies which sum to 1. Patches can exist in two states, either saturated or empty. Saturated patches are occupied by hosts that dominate the local resources, and cannot be invaded. Patch clearing events, which occur with rate *c* per patch per unit time, empty saturated patches of their resident hosts, and enable invasion by strains from other patches. Invasions of empty patches occur with rate *m* per patch per unit time, which specifies the influx rate into a patch. We will assume that empty patches are rapidly invaded, i.e. *c* ≪ *m*, and that patch clearing is the rate limiting step to initial patch invasion. When invasions occur, they bring in a small inoculum from a patch of type *i* with probability *x*_*i*_. Patch clearing and migration mimic natural turnover events that occur e.g. in the gut due to peristalsis, in soil microenvironments due to physical perturbations, or during host-to-host transmission in epidemiology.

In the initial stages of patch invasion, we postulate that it takes a certain establishment time, *τ*_*est*_, for an invading strain to grow sufficiently to saturate the patch and thereafter exclude other invaders. If no further migration events occur during time *τ*_*est*_ after the initial invasion, i.e. *τ*_*est*_ ≪ 1/*m*, the patch will become occupied and dominated by the invading type *i*. The probability of such single invasion events is *e*^−*mτ*_*est*_^ ≈ 1 − *mτ*_*est*_, and the total rate of such events is *c*(1−*mτ*_*est*_) ≈ *c*. Since the patch clearing rate is independent of patch type, and invading strains are chosen randomly according to the distribution *x*_*i*_, there will be no net change of *x*_*i*_ due to single strain invasion events. Thus, the dynamics of *x*_*i*_ are determined by multi-strain invasion events, which occur with total rate 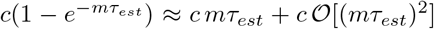. We can therefore disregard invasion events with three or more strains if *mτ*_*est*_ ≪ 1, e.g. for short establishment time and/or rare immigration, and consider the dynamics due to co-invasion events, i.e. with exactly two invading strains in competition.

Co-invasion events by strains of types *i* and *j* occur with rate 2*c mτ*_*est*_*x*_*i*_*x*_*j*_, for *i* ≠ *j*, and rate 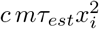 for *i* = *j*. If *i* = *j*, the types are identical, and the patch becomes saturated with type *i*. If *i* ≠ *j*, competition occurs and resolves itself over a timescale that is determined by the fitness difference *f*_*ij*_ ≡ |*f*_*i*_ − *f*_*j*_| between the two types, where *f*_*i*_ is the fitness of type *i*. When *f*_*i*_ > *f*_*j*_, the competition will resolve in favor of type *i*, provided that the competition is over before the next patch clearing event. If patch clearing occurs before resolution, then neither type makes any gains. The probability of successful resolution in favor of type *i* is thus *p*_*ij*_ ≡ *f*_*ij*_/(*f*_*ij*_ + *c*), if *f*_*i*_ > *f*_*j*_, or *p*_*ij*_ = 0 if *f*_*i*_ < *f*_*j*_. The total rate of co-invasions of *i* ≠ *j* successfully resolving in favor of *i* is 2*c mτ*_*est*_*p*_*ij*_*x*_*i*_*x*_*j*_. The replicator equation for this system is

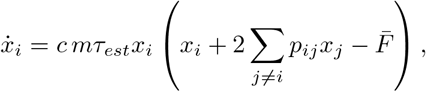

where

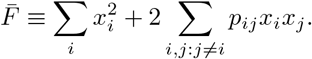

Note that resolution of the competition increases the number of patches occupied by the winning type, but it does not decrease the number of patches occupied by the loser; rather, it decreases the *frequency* of all other patch types. In other words, this is not a zero sum game.

Since invasion events bring a representative inoculum from a saturated patch, invasion by **R**_*s*_ will transfer both the resistance switching strain and the phage while **R**_0_ and **S** transfer host cells of a single phenotype. For a preventative defense, the competition between **R**_0_ and **R**_*s*_ will result in the former driving both the latter strain and the phage to extinction, at rate 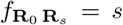. In a competition of **R**_*s*_ versus **S**, the phage will rapidly drive strain **S** to extinction, at rate 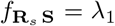, while competition of **S** versus **R**_0_ results in the former outcompeting the latter at rate 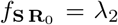 due to the cost of defense in the absence of phage (Fig. 4A); we do not require the exact expressions of *λ*_1_ and *λ*_2_, and only assume that *λ*_1_, *λ*_2_ > *c*. The invasion probabilities *p*_*ij*_ will then reflect fitness differences shown in Fig. 5A, and the payoff matrix for this game is given by

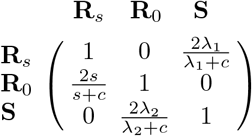

**FIG. 5.**
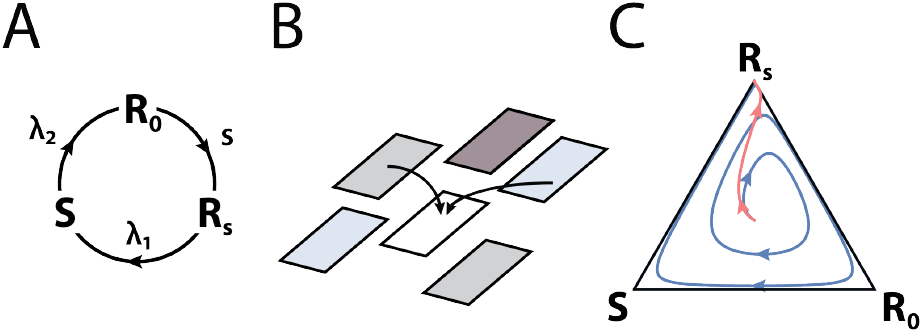
Patch invasion dynamics. (A) Illustration of the rock-paper-scissors-type dynamics among a resistance switching strain **R**_*s*_ (which carries phage), a non-switching strain **R**_0_, and a sensitive strain **S**. **R**_0_ beats **R**_*s*_ with rate *s*, **S** beats **R**_0_ with rate λ_2_, and **R**_*s*_ beats **S** with rate λ_1_. (B) The patch invasion model considers co-invasion events into a cleared patch. Patch clearing events occur at rate *c*. Shading reflects different bacterial strains. (C) Patch invasion game dynamics approaches a heteroclinic cycle on the boundary for *s* > *c* (blue curve) or the strict Nash equilibrium **R**_*s*_ for *s* < *c* (pink curve).

It is now clear that for *s* > *c* this is a rock-paper-scissors game with no stable Nash equilibria and a heteroclinic cycle on the boundary (Fig. 5C, blue curve) [35]. However, for *s* < *c* a strict Nash equilibrium (i.e. an ESS) exists, which corresponds to the resistance switching strain **R**_*s*_ (Fig. 5C, pink curve).

We can further consider a set of **R**_*s*_ strains spanning a wide range of values of *s*, either pre-existing or occurring by mutation (*Supplementary Materials*). Competitions between strains **R**_*s*_ and **R**_*s*′_ will resolve in favor of the strain with a smaller switching rate, provided that |*s* − *s*′| > *c*. Thus, the distribution of switching rates across the ecology will evolve toward lower values of *s* until all remaining **R**_*s*_ strains satisfy *s* < *c*, rendering **R**_0_ unable to invade. At that point, any remaining **R**_0_ and **S** strains will be driven to extinction, and resistance will be preserved thereafter across the ecology.

Finally, we comment on the patch invasion dynamics for immune defenses. The **R**_**0**_ strain can generate **R**_0_-*SP* coexistence exclusively through co-invasions with the **R**_**s**_ strain, and can no longer be invaded by the **S** strain. The ecology for immune defenses will therefore contain patches with an **R**_0_-*SP* coexistence, as well as patches occupied by **R**_*s*_ strains with *s* < *c*.

## II. DISCUSSION

By analyzing host-pathogen dynamics within a single patch and modeling a minimal ecology consisting of many patches, we determined the set of conditions in which resistance mechanisms are preserved. We showed that the resistance switching strategy, whereby pathogen-resistant hosts stochastically lose resistance, enables the ecology as a whole to maintain memory of the pathogen. Such *ecological memory*, an emergent property at the level of ecological dynamics, is the basic requirement for preserving resistance mechanisms over long timescales. Our analysis shows that a non-zero failure rate of preventative defense mechanisms, which reduces host growth rate within a single patch, protects those same mechanisms from eventual loss across the ecology of patches.

Spontaneous loss of a preventative defense was observed experimentally to enable persistence of phage at low frequencies [10, 23, 24, 36]. Similarly, loss of immune defenses is known to occur in CRISPR systems [34, 37, 38], and modeling has shown that this loss could be responsible for coexistence of phage and bacteria [39], which was experimentally observed in [40]. Our work unifies these observations by providing the critical context of ecological dynamics and memory, and thereby establishes a rigorous basis to analyze evolutionary maintenance of resistance mechanisms. We showed that the same resistance switching strategy that enables ecological memory can maintain multiple phage strains with different combinations of infection rates and burst sizes. Ecological memory can therefore involve a diverse collection of phage types that coexist stably with the host bacterium, which has implications for microbial ecosystem diversity and stability. Further extension of our modeling approach for immune defenses that accounts for CRISPR spacer acquisition and evasion by phage [41–45] may be fruitful in identifying novel dynamics and strategies of generation and maintenance of ecological memory.

We showed that the specific molecular mechanism of resistance has a large effect on host-pathogen dynamics. In contrast to preventative defenses, immune mechanisms act as phage sinks, absorbing and removing phage from the environment. Phage-sensitive strains, which do not pay the cost of immunity, can exploit immune strains to enable their own survival in the presence of phage. On the one hand, this means that immune defenses cannot be eliminated by faster-growing sensitive strains, as these depend on the mutualism for survival. On the other hand, cheating reduces the long-term growth potential of immune strains and may thus select for anti-cheating strategies. We found that resistance switching by immune strains increases the amount of phage in the environment and can be used to select against cheater strains.

The models introduced here enable testing and validation in laboratory experiments. Our single patch formulation corresponds to a well-mixed population of bacteria and phage growing in rich or limited media and maintained in a proliferating state by dilution. The host-pathogen interaction term, *k*_*I*_ (*t*), was constructed by considering phage-receptor binding interactions, yielding a general form applicable across different regimes of host and phage densities, spanning different experimental scenarios. A major prediction of our single-patch models is that immune and sensitive strains can coexist stably with phage, which can be directly tested by growing mixtures of bacterial strains with and without a CRISPR system in the presence of phage. Depending on the phage burst and infection rate parameters, our model predicts whether or not coexistence is possible; comparison with a preventative defense in the same strain (e.g. a phage receptor loss mutant), where coexistence is predicted not to occur, would yield further validation. Our multi-patch ecological model can be tested in multi-well plate format experiments using the *λ*-phage system, where each well corresponds to a patch and is inoculated with one of three possible *E.coli* strains, **R**_0_ (*lamB* deletion), **S** (constitutive *lamB*), and **R**_*s*_ (wild type) with phage, in media and grown to saturation. Daily dilution into fresh plates would be performed such that each well receives inocula from two randomly chosen wells of the saturated culture. Ecological dynamics are observed by tracking the prevalence of resistance across the plate.

While our modeling considered phage-bacteria interactions, the general principles that we identified are likely to be relevant in a wide range of systems, including vertebrate immune systems [46, 47] and other epidemiological dynamics [48]. In particular, our formulation of patch invasion dynamics using game theory, together with the mechanism of ecological memory, may be applicable to the maintenance of pathogen resistance mechanisms in plants, as the costs of such resistance are well-known and patch dynamics models are widely used in plant ecology [3, 49, 50]. As observed in [23, 36], resistance switching in bacteria not only sustains the growth of phage but also allows the pathogen eventually to evolve to infect the host through a different pathway. We propose that ecological memory is a general mechanism that enables the long-term co-evolution of pathogens and hosts.

## ACKNOWLEDGMENTS

We thank David Gresham, Enrique Rojas, Mark Siegal and Joshua Weitz for discussions. This work was funded by NIH grant R01-GM120231 to E.K.

## Appendix A: Host infection rate *k*_*I*_ (*t*)

We consider a phage-receptor binding reaction

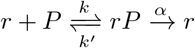

where *r* denotes a receptor and *rP* a phage-receptor complex. The reversible binding/unbinding of phage occurs at rates *k* and *k*′ whereas the absorption of a bound phage occurs with rate *α* [27–29]. In a quasi-steady state approximation the concentration of the complex *rP* is constant over timescales 1/*α* which can be very short, and we solve the quadratic equation for *rP* to obtain

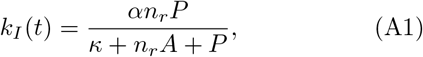

where *n*_*r*_ is the average number of receptors per host, and *κ* = (*k*′ + *α*)*V*/*k* is the Michaelis constant for phage-receptor kinetics. This form of interaction enables studying a range of systems, from high phage-receptor binding affinities (*κ* → 0) to low binding affinities. For *κ* ≫ *P, n*_*r*_*A* we recover the typical Lotka-Volterra interaction term which we analyze using chemostat control (*Supplementary Materials*, Fig. S2). The results presented in the paper consider minimal sensitivity, given with *n*_*r*_ = 1, to identify a minimal condition for coexistence, while in *Supplementary Materials* we consider hosts with any number of receptors.

## Appendix B: Fixed points and bifurcation curves

The system of equations (1) can exhibit unbounded growth, which can be limited by additional terms, e.g. by including turbidostat or chemostat control. We obtain the analytic form of bifurcation curves by linear stability analysis on the fixed points of the bounded system. For turbidostat control, we obtain:

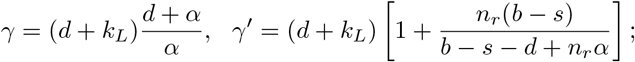

similar expressions exist for chemostat control (*Supplementary Materials*). The curve *γ*″ is found numerically (*Supplementary Materials*).

## Appendix C: Host extinction stability

For a host with the resistant phenotype the extinction fixed point E is never stable. In the absence of resistance we perform the stability analysis on a reduced system for which *R* = 0. For *α* < *d*/*n*_*r*_, E is unstable. For *α* > *d*/*n*_*r*_, E is stable at any *βk*_*L*_, and since S is stable for *βk*_*L*_ < *γ*, we obtain the bistable region S&E shown in Figs. 2E and 3C.

## Supplementary Material

### S1. THE MODEL OF PHAGE ABSORPTION

To derive the functional form of the host infection rate we consider a model of phage absorption at a molecular level where a phage reversibly attaches to a receptor followed by injection of its genetic material. The kinetics are given with

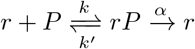

The receptor *r* reversibly binds/unbinds a phage *P* at rates *k* and *k*′, forming an intermediate phage-receptor complex *rP*. The rate *α* denotes the phage absorption rate by the receptor. In this reaction, the quantity of *r* is preserved and phage is removed. Phage compete for all the receptors, and a host, which can carry multiple receptors, will bind multiple phage. In the quasi steady-state approximation the concentration of phage-receptor complex, [*rP*], satisfies

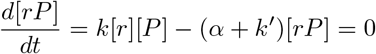

By writing [*r*] = [*r*_*tot*_] − [*rP*] and [*P*] = [*P*_*tot*_] − [*rP*] for free receptor and phage concentrations we obtain a quadratic equation for [*rP*]

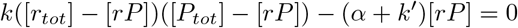

which is solved with

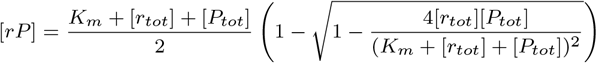

where *K*_*m*_ = (*k*′ + *α*)/*k* is the Michaelis constant. This equation is known as the Morrison equation. Since 4[*r*_*tot*_][*P*_*tot*_]/(*K*_*m*_ + [*r*_*tot*_] + [*P*_*tot*_])^2^ < 1 we expand the square root in a power series and stop at the first order to obtain:

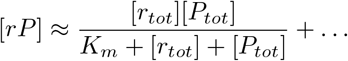

where corrections are of the form [*r*_*tot*_]^*n*^[*P*_*tot*_]^*n*^/(*K*_*m*_ + [*r*_*tot*_] + [*P*_*tot*_])^2*n*−1^ for *n* = 2, 3, …. By setting [*r*_*tot*_] = *n*_*r*_[*A*_*tot*_] where *n*_*r*_ is the average number of receptors per cell, we obtain the total phage absorption rate

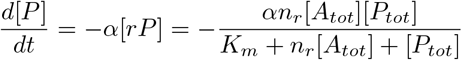

Once the receptor on the sensitive cell absorbs a phage, the host becomes infected and acts as a sink for phage that subsequently bind to its receptors. Since [*A*_*tot*_] = [*S*_*tot*_]+[*I*_*tot*_] for preventative defenses (or [*A*_*tot*_] = [*R*_*tot*_]+ [*S*_*tot*_] + [*I*_*tot*_] for immune defenses), the infection rate, which corresponds to a part of this process that converts *S* to *I*, is equal to

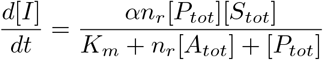

From here we obtain the infection rate *k*_*I*_ provided in the main text by multiplying the numerator and denominator by system volume, which converts concentrations to population size

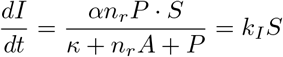

where the constant κ = *K*_*m*_*V* sets the binding affinity, with respect to which different limits can be achieved:

- *the low binding affnity limit κ* ≫ *n*_*r*_*A* leads to the Michaelis-Menten rate

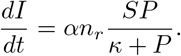
- *the Lotka-Volterra limit κ* ≫ *n*_*r*_*A*, *P* obtains

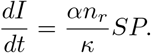
- *the high binding affinity limit n*_*r*_*A* ≫ *κ* in which we omit *κ* to obtain

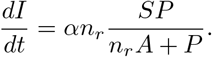

### S2. ANALYSIS OF THE MODEL OF PREVENTATIVE OR IMMUNE DEFENSE

#### S2.1. Projection onto a simplex

To analyze the dynamics of population structure, we compute the relative biomass fractions of each subpopulation within the total system volume *V* . The total biomass is given by *B* ≡ *ν*_*H*_ (*R* + *S* + *I*) + *ν*_*P*_ *P*, where *ν*_*H*_ and *ν*_*P*_ are the average mass of a host cell and a phage, respectively. The biomass fractions are given by *f*_*R*_ = *ν*_*H*_ *R*/*B*, *f*_*S*_ = *ν*_*H*_ *S*/*B*, *f*_*I*_ = *ν*_*H*_ *I*/*B*, and *f*_*P*_ = *ν*_*P*_ *P*/*B*; and since *f*_*R*_ + *f*_*S*_ + *f*_*I*_ + *f*_*P*_ = 1, the biomass fractions specify a point on the simplex. The host infection rate *k*_*I*_ (*t*) is expressed as

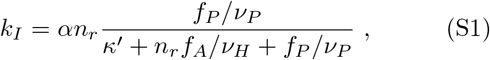

where *f*_*A*_ = *ν_H_ A/B* is the total biomass fraction of phage-absorbing hosts, and *κ*′ = *K_m_V/B* is the binding constant divided by the fixed total biomass density *B/V* . In the high binding affinity limit, which we analyze here, we omit *κ*′; we comment in a section below how inclusion of *κ*′ modifies the stability of fixed points. The population dynamics equations on the simplex are given by

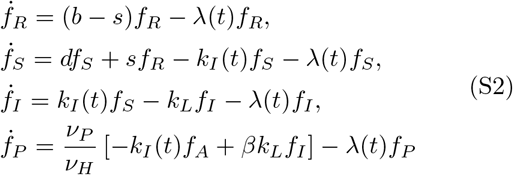

where 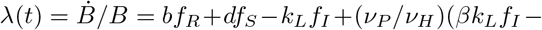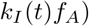 is the instantaneous biomass growth rate. The equations depend on the ratio of phage to host biomass *ν* ≡ *ν*_*p*_/*ν*_*H*_. As we will show, the stability of the fixed points, which determine the phase structure, does not depend on the choice of *ν* in the high binding affinity limit. Additionally, we show in section S2 S2.7 that turbidostat control of host biomass yields equations that can be mapped by a smooth, 1-to-1 mapping to the equations above; the growth rate and linear stability analysis in a turbidostat therefore match those obtained at the corresponding fixed points on the simplex.

#### S2.2. Linear stability analysis

We perform linear stability analysis of the fixed points of (S2), by computing the eigenvalues of the Jacobian matrix, whose elements are 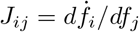. Since Σ_*i*_ *f*_*i*_ = 1, we express *f*_*P*_ = 1 − *f*_*R*_ − *f*_*S*_ − *f*_*I*_ and solve a reduced system consisting of the first three equations in (S2).

##### E phase

The host extinction fixed point E corresponds to the solution {*f*_*R*_, *f*_*S*_, *f*_*I*_} = {0, 0, 0}. The eigenvalues of the Jacobian evaluated at E are

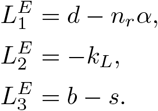

Since 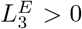, extinction is always unstable in the presence of resistant phenotype. Analysis of host extinction stability for a system with *f*_*R*_ = 0 is presented further below.

##### S phase

The S phase corresponds to the solution {*f*_*R*_, *f*_*S*_, *f*_*I*_} = {0, 1, 0}. Eigenvalues of the Jacobian matrix are

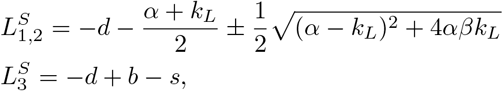

which are negative for *d* > *b* − *s* and

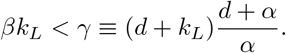

##### SP phase

Similarly we analyze the SP phase which corresponds to the solution 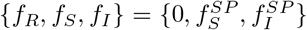. One of the eigenvalues of the Jacobian evaluated at the SP fixed point is

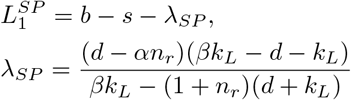

where *λ*_*SP*_ is the growth rate in this phase, while the other two eigenvalues are both negative for *α* < *d*/*n*_*r*_ and *βk*_*L*_ > *γ*. Solving the 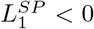 condition for *βk*_*L*_ in the region *α* < *d*/*n*_*r*_ and *βk*_*L*_ > *γ* gives the location of the stable SP phase:

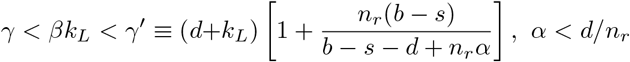

##### RSP phase

So far, the eigenvalues were real and the analysis holds for both preventative and immune defenses. For *βk*_*L*_ > *γ*′ the 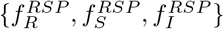 fixed point exists and its stability is determined by considering eigenvalues which can also be in the complex domain. The RSP phase can therefore contain stable fixed points and periodic dynamics that emerge in the regions where there are no stable fixed points. The existence and size of these regions in the phase diagram will depend on which of the two models of resistance we consider. To find the location of the curve *γ*″ where complex-conjugate eigenvalues become purely imaginary, and therefore can lead to a Hopf bifurcation of the dynamical system, we consider the characteristic equation, which is a cubic polynomial in eigenvalues *L* with real coefficients *a*_*i*_: *P* (*L*) = *L*^3^ + *a*_2_*L*^2^ + *a*_1_*L* + *a*_0_ = 0. The solutions will satisfy Viete’s formulas *L*_1_ + *L*_2_ + *L*_3_ = −*a*_2_, *L*_1_(*L*_2_ + *L*_3_)+ *L*_2_*L*_3_ = *a*_1_, and *L*_1_*L*_2_*L*_3_ = −*a*_0_. Let *L*_1_ be the real eigenvalue, which is negative when *a*_0_ > 0, and let *L*_2_ and *L*_3_ be the complex conjugate eigenvalues. The fixed point will become unstable when the real parts of *L*_2_ and *L*_3_ vanish, which when used with Viete’s formulas gives a condition *a*_1_*a*_2_ = *a*_0_ that we solve for *βk*_*L*_.

#### S2.3. Phase diagram without resistance

Now we consider Eq. (S2) with *f*_*R*_ = 0, corresponding to a host which has not acquired resistance. This reduces the system to two equations for *f*_*S*_ and *f*_*I*_ , and the eigenvalues of the Jacobian are computed by solving a minor of the full model with rows and columns corresponding to the resistant phenotype removed. Figure S1 shows the phase diagram displaying the regions of stable S, SP and E fixed points in the absence of resistant phenotype.

The eigenvalues at the host extinction fixed point {*f*_*S*_, *f*_*I*_} = {0, 0} are

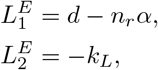

which make extinction stable for *α* > *d*/*n*_*r*_. The S phase {*f*_*S*_, *f*_*I*_} = {1, 0} is stable for *βk*_*L*_ < *γ*. Thus, there is a bistable region at *α* > *d*/*n*_*r*_ and *βk*_*L*_ < *γ* which is shown in Fig. S1. The SP phase is stable for *α* < *d*/*n*_*r*_ and *βk*_*L*_ > *γ*. In this reduced model, in the region where E is stable there can exist a second extinction fixed point, such that *f*_*S*_ = 0 and *f*_*I*_ = const > 0, and in which infected host and phage decay exponentially. This pathological fixed point is never stable in the full model with resistance, but in the reduced model we consider it part of the E phases.

#### S2.4. Plotting flow diagrams

The choice of host and phage mass units in *ν* = *ν*_*P*_/*ν*_*H*_ has no effect on stability of the phases, since bifurcation curves are determined by the eigenvalues of the Jacobian and are therefore basis independent. These analyses generally hold for any host-phage system whose dynamics are described with Eq. (S2). To plot the flow diagrams in Figs. 2A-D & 3A-B in the main text, we set *ν* = 1, and therefore show the relative abundances of host and phage in a population. Points in the interior of the diagram can be read by following the dotted gridlines to each edge, as illustrated in this example:

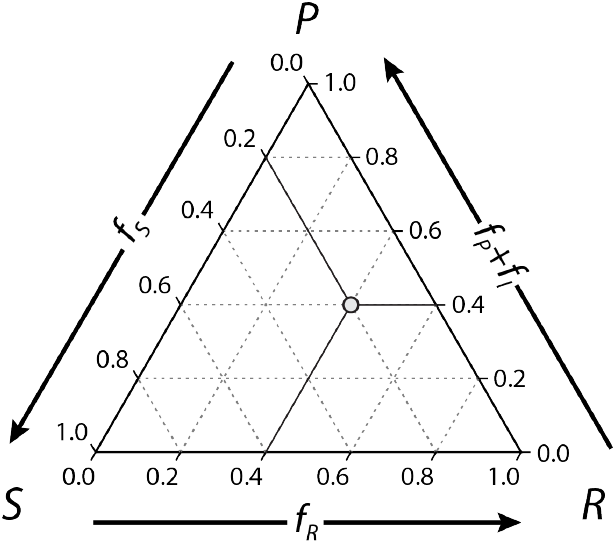

#### S2.5. Growth rate

From the first equation in (S2) we observe that when the RSP fixed point is stable, the system growth rate is *λ* = *b − s*. For stable fixed points with *f*_*R*_ = 0, we use *ḟ*_*S*_ = 0 to obtain *λ* = *d* in the S phase and *λ* = (*d* − *αn*_*r*_)(*βk*_*L*_ − *d* − *k*_*L*_)/(*βk*_*L*_ − (1 + *n*_*r*_)(*d* + *k*_*L*_)) in the SP phase. For periodic dynamics, at steady state the average growth rate can be obtained by integrating *ḟ*_*R*_/*f*_*R*_ over the period *T*:

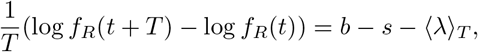

where 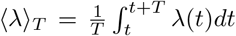. Since *f*_*R*_(*t* + *T*) = *f*_*R*_(*t*), we obtain ⟨*λ*⟩ _*T*_ = *b − s*, the growth rate of the resistant phenotype.

#### S2.6. Stability analysis at low binding affinity

We examine the stability of phases for *κ*′ > 0 by considering its modifications to the host infection rate.

##### Immune defenses

For immune defenses, where *f*_*A*_ = *f*_*R*_ + *f*_*S*_ + *f*_*I*_ , the infection rate for *κ*′ > 0, given with (S1), can be rewritten as

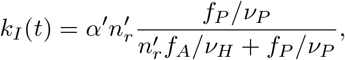

where we introduced rescaled parameters 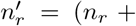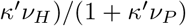 and *α*′ = *αn*_*r*_/(*n*_*r*_ + *κ*′*ν*_*H*_). Since this expression matches the functional form of the infection rate in the high binding affinity limit, the phase diagram of immune defenses for *κ*′ > 0 maintains the structure shown in the high binding affinity limit, with replacements *α* ↔ *α*′ and 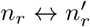.

##### Preventative defenses

For preventative defenses we obtain

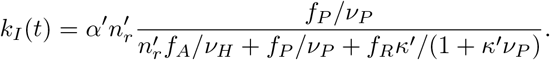

The linear stability analysis of the S and SP phases where *f*_*R*_ = 0 recovers the same results as for the immune defenses with *α*′ and 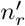, while the RSP phase could potentially include limit cycles.

#### S2.7. Turbidostat control of host biomass

We consider the control of host biomass by a feedback dilution mechanism that maintains a fixed host density in the system volume *V* . The dynamics are given by transforming the simplex equations (S2) to a set of coordinates given by phenotype frequencies in the host population *x* = *R/*(*R* + *S* + *I*), *y* = *S/*(*R* + *S* + *I*), *z* = *I/*(*R* + *S* + *I*) = 1 − *x* − *y* and phage pressure *p* = *P/*(*R* + *S* + *I*). In this basis, the infection rate is *k*_*I*_ = *αn*_*r*_*p*/(*κ*_*H*_ + *n*_*r*_*a* + *p*), with *κ*_*H*_ ≡ *K*_*m*_*V*/(*R* + *S* + *I*) and where *a* = *A/*(*R* + *S* + *I*) is total frequency of phage-adsorbing hosts, which equals 1 − *x* for preventative and 1 for immune defenses. The turbidostat equations are:

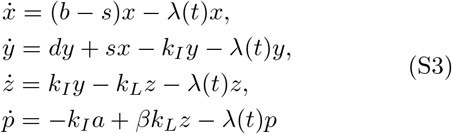

where *λ*(*t*) = *dy* + *bx* − *k*_*L*_*z* is the instantaneous host growth rate. These coordinates can be mapped to the simplex via:

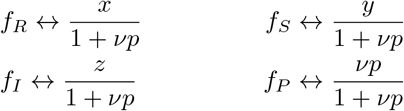

except at the host extinction point E, which is a singular outcome in this basis and needs to be studied on the simplex. The stability of fixed points in the host biomass basis matches that on the simplex and generates the same phases as the ones presented for the simplex equations.

#### S2.8. Chemostat control of host biomass

We analyze the dynamics in a chemostat model given by the equations

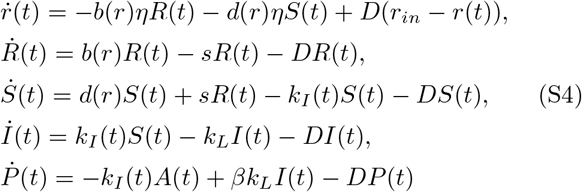

Here *r* is the resource, *η* is the resource to growth conversion factor (which we set to 1), *r*_*in*_ is the concentration of the supplied resource while *D* is the chemostat dilution rate. *R*, *S*, *I* and *P* denote concentrations of host and phage, and the infection rate is *k*_*I*_ = *αn*_*r*_*P*/(*K*_*m*_ + *n*_*r*_*A* + *P*). We consider the high binding affinity limit and set *K*_*m*_ = 0. Cellular division rates depend on resource concentration and are generally chosen to be Monod functions of the form *d*(*r*) = *d r/*(*M* + *r*), where *M* is nutrient concentration at half-velocity.

As exponential growth is achieved in a condition of resource saturation *r* ≫ *M*, here will we will explore the regime of nutrient limitation *r* ≪ *M*, for which cellular division rates linearly depend on *r*: *d*(*r*) = *d·r*/*M*. Hereafter we set *M* = 1. Extinction of both phage and host is globally stable if *D* > *d · r*_*in*_, which means the population is washed out faster than it can grow. Dilution rates below *d · r*_*in*_ but above *b · r*_*in*_ will select against the resistant phenotype. We will consider dilution rates *D* < *b · r*_*in*_ − *s* so that the chemostat supports growth of both resistance-switching and sensitive phenotypes.

The analysis of chemostat fixed points gives the same phases as in the turbidostat formulation with: (i) the *E* phase, which is unstable for our choice of dilution rate; the *S* phase, corresponding to the solution *r* = *D*/*d*, *S* = *r*_*in*_ − *r*, *R* = *L* = *P* = 0, which is stable for

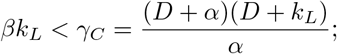

(ii) the *SP* phase which is stable for 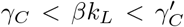 where

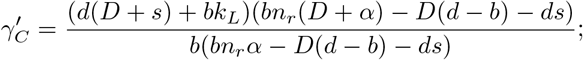

and (iv) the *RSP* phase, whose stability we will determine numerically. Since the dilution rate controls the overall growth rate and therefore the relative fitness differences between phenotypes, changing the dilution rate will impact the stability of the fixed points and can lead to removal or enhancement of regions that admit periodic dynamics in the phase diagram.

Figure S2 shows the representative phase diagrams of stable fixed points in the chemostat model for preventative and immune defenses, and in the Lotka-Volterra limit where the infection rates are *k*_*I*_ = *k*_*LV*_ *P* with *k*_*LV*_ = *αn*_*r*_/*K*_*m*_.

#### S2.9. Transcritical bifurcations

Figure S3 plots the growth rate and phenotype and phage frequencies as a function of *α* for fixed *β*, showing that as *α* increases from the S to SP to RSP phase, the system goes through two transcritical bifurcations.

#### S2.10. Bistable regions RSP&S in the preventative defense model

Figure S4 shows additional phase diagrams featuring RSP&S bistable regions, that we obtain for large switching rates and/or large costs of defense, which are given with *d* − *b*.

#### S2.11. Switching maintains multiple phage types in the preventative defense model

Here we consider *P*_1_ and *P*_2_ that compete for the same host. In this model, a host cannot be simultaneously infected by more than one phage type, but the infected cell can absorb phage of any type. The two phage types can have different absorption rates *α* and burst sizes *β* and we ask under which conditions they coexist. We model cell infection rate as in section S1, which provides the 2-phage form of the infection rate shown in the equations below:

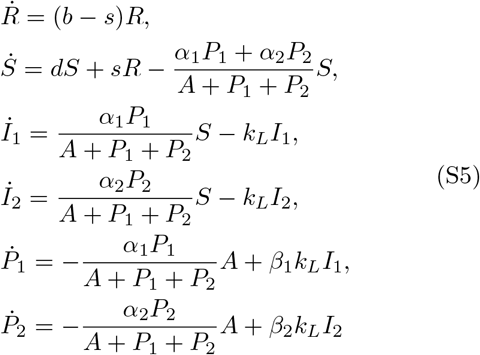

where *A* = *S* + *I*_1_ + *I*_2_. Here we consider the minimal sensitivity condition *n*_*r*_ = 1, but the model is easily generalizable for any *n_r_ >* 1. Figure S5 shows the phase diagrams for *P*_2_ given *P*_1_ with coordinates {*α*_1_*, β*_1_} at three different locations in the RSP phase. Dashed curves denote transitions from single to multiple phage phases where both phage types are maintained in the RSP phase. In the region where *P*_2_ outcompetes *P*_1_ resistance can be lost if *P*_2_ parameters are located in *P*_2_’s *SP* phase; this region occurs at high *P*_2_ burst rates and is shown in Fig. S5 in pink. The size and shape of these regions does not strongly depend on the value of *s* > 0. For zero switching rate there will be no phage coexistence, since the RSP phase collapses to pure R, which does not support phage growth.

#### S2.12. Varying lysing rates does not change the phase structure

Figure S6 shows phase diagrams as a function of the lysing rate *k*_*L*_. We varied *k*_*L*_ by a factor of two above and below cell division rate *d* to show that the phase structure remains unchanged over the range that is commonly found in experiments. Larger deviations of *k*_*L*_ similarly do not affect the phase structure.

#### S2.13. Phase diagrams for a large number of receptors per cell

We consider the phase structure of Eq. (S2) for a range of *n*_*r*_ including *n*_*r*_ ≫ 1. In the limit *n*_*r*_ → ∞ the dependence on *n*_*r*_ drops out, as

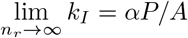

Figure S7 shows the phase diagrams for *n*_*r*_ = 1, 100 and ∞, for preventative and immune defenses. As *n*_*r*_ increases the SP phase moves to lower values of *α* < *d*/*n*_*r*_ while the RSP phase for immune defenses develops a larger region where periodic dynamics are possible. In the limit *n*_*r*_ → ∞ the SP phase disappears and the phase diagram contains only the S and RSP phases separated with *γ* = (*d* + *α*)(*d* + *k*_*L*_)/*α*.

#### S2.14. Phase diagrams for different costs of defense

Figure S8 shows phase diagrams as a function of the resistant phenotype division rate *b*. The cost of the defense is reflected in the difference between the division rates of sensitive and resistant cells, *d* − *b*. For larger costs, the SP phase becomes wider for both defenses. For immune defenses the height of the Hopf bifurcation curve increases, while the changes in stability of the RSP phase for preventative defenses are shown in Fig. S4.

#### S2.15. Phase diagrams for the case where phage stays bound to the receptor

We also consider phage that does not fall off the receptor upon infection and therefore blocks the receptor for subsequent absorption of phage in the infected phenotype. This effect would be small for large *n*_*r*_ and most extreme in the model of minimal sensitivity (*n*_*r*_ = 1), which would prevent the infected phenotype to act as a phage sink. We solved the *n*_*r*_ = 1 model to find this change shifts the location of *γ*′ and *γ*″ bifurcations as well as impacts the dynamics at the intersection of S, SP and RSP phases. Figure S9 shows the phase diagrams for preventative and immune defenses.

#### S2.16. Periodic orbits in the model of immune defense

Figure S10 shows an example of a contraction of the periodic orbits with the increase in *s*. For small switching rates the orbit passes close to the boundary of the simplex; as *s* is increased the orbit contracts towards the unstable fixed point in the interior, which for a critical switching rate *s*^∗^ ≈ 0.0216 becomes stable.

#### S2.17. Phage decay

Here we consider the effects of introducing a phage decay rate *δ* in our model, which now becomes

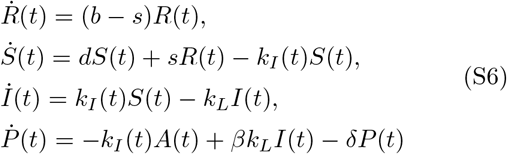

We analyze the fixed points as before to obtain the formulae for bifurcation curves:

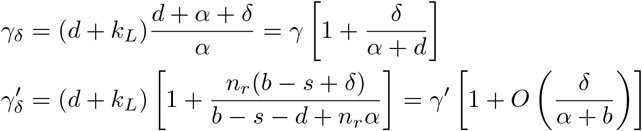

Since phage decays typically on the order of days [1], such corrections are small and only slightly shift bifurcation curves.

#### S2.18. Model of CRISPR spacer loss

We modify the immune defense model to account for a loss-of-spacer phenotype, *S*, which is sensitive to phage but pays the cost of expressing the immune system (i.e. it grows at the same rate as *R*), and which occurs by switching from *R* at a spacer loss rate *s*. We examine the coexistence of the resistant strain with a cheater strain *S*′ which is sensitive to phage and grows at rate *d*′ > *b*. The dynamics are given with:

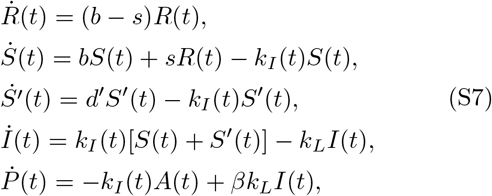

where *A* = *R* + *S* + *S*′ + *I*. In this system there exist five stable fixed points that determine late-time population structure: the previously described S, SP and E phases, and two new phases that carry resistance, the S’RSP phase where all hosts coexist and the cheater mooches off the resident CRISPR defense, and the RSP phase where the cheater is removed. Figure S11 shows the diagram of these phases separated by curves across which the system undergoes transcritical (*γ*, *γ*′, *γ*‴) and Hopf (*γ*″) bifurcations.

Starting from a point in the S’RSP phase and increasing phage burst rate, the frequency of *S*′ decreases until it becomes exactly zero at the location of the curve *γ*‴. The dynamics transitions to a stable fixed point where the cheater is completely cleared from the patch. Stability of the RSP phase implies that any transient increases of *S*′ frequency, e.g. through random mutations or immigration events, will decay exponentially. The location of this transition is controlled by the fitness difference between *S*′ and *S* which includes the cost of resistance, and the rate of spacer loss:

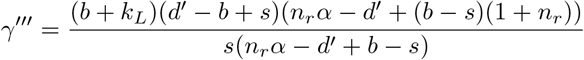

For *α* ≫ *b* we obtain the lower bound on burst size for which the cheater is removed:

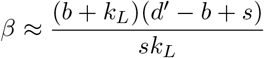

Therefore, for *βs* ≳ Δ*f*, where Δ*f* = *d*′ − *b* + *s* is the fitness difference between the resistant switcher strain and the cheater, loss of resistance suppresses invasions by *S*′. Increasing the switching rate increases the selection against cheating.

Note that for simplicity we considered only spacer loss and not CRISPR loss; generally, both spacer loss and CRISPR loss will be present, further increasing the selection to remove the invader from the patch.

### S3. PATCH INVASION MODEL

#### S3.1. Generalization of the patch invasion model

Here we discuss the game theory model given with a payoff matrix:

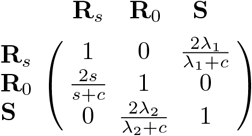

Since we can subtract a constant 1 from each column without changing the equilibria, we have equivalently

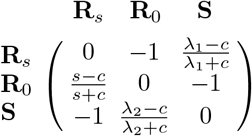

From here it becomes clear that for *s* < *c* strain **R**_0_ no longer beats **R**_*s*_ and a strict Nash equilibrium exists which corresponds to the pure strategy **R**_*s*_.

Now we consider a model with two switching strains, **R**_*s*_ and **R**_*s*′_, with *s − s*′ > *c*. In a similar fashion, we obtain the payoff matrix *ϕ*:

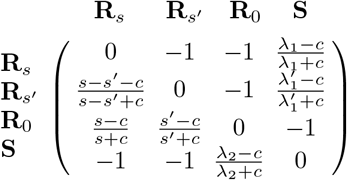

We analyze the fixed points of the replicator equation corresponding to matrix *ϕ*,

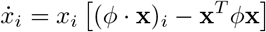

where *i* = {**R**_*s*_, **R**_*s*′_, **R**_0_, **S**}. The stable fixed point corresponds to a pure 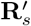 strategy for *s*′ *< c*. If both *s* and *s*′ are larger than *c*, there is no interior equilibrium and the system will approach the boundary of the tetrahedron Σ_*i*_ *x*_*i*_ = 1 whose faces correspond to a reduction of the game theory model to a subset of three strains. The unstable fixed point on the {**R**_*s*_, **R**_0_, **S**} face repels interior orbits and the system transitions to a heteroclinic cycle on the {**R**_*s*′_, **R**_0_, **S**} face. Equivalently, the strain with the highest switching rate is driven to extinction as it gets displaced by the strain with lower switching rate and the ecology reduces to the {**R**_*s*′_, **R**_0_, **S**} patch invasion game.

Now we can generalize to a large number of switching strains with *s* > *s*′ > *s*″ > … > *c* where the difference between each pair of switching rates is greater than *c*. The invasion diagram for three switching strains is:

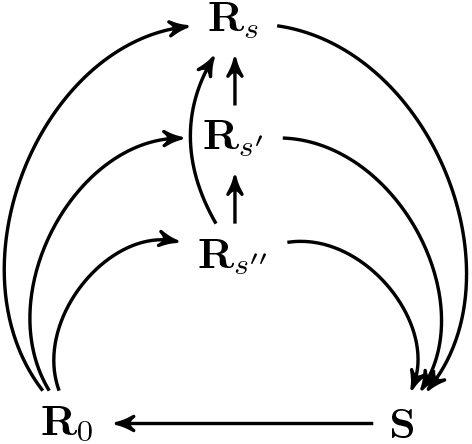

If all the strains have switching rates above *c*, the ecology will undergo a similar reduction where the strains with highest switching rates progressively go extinct until one of the rates evolves to become smaller than *c*, at which point it becomes an ESS.

---

**FIG. S1.**
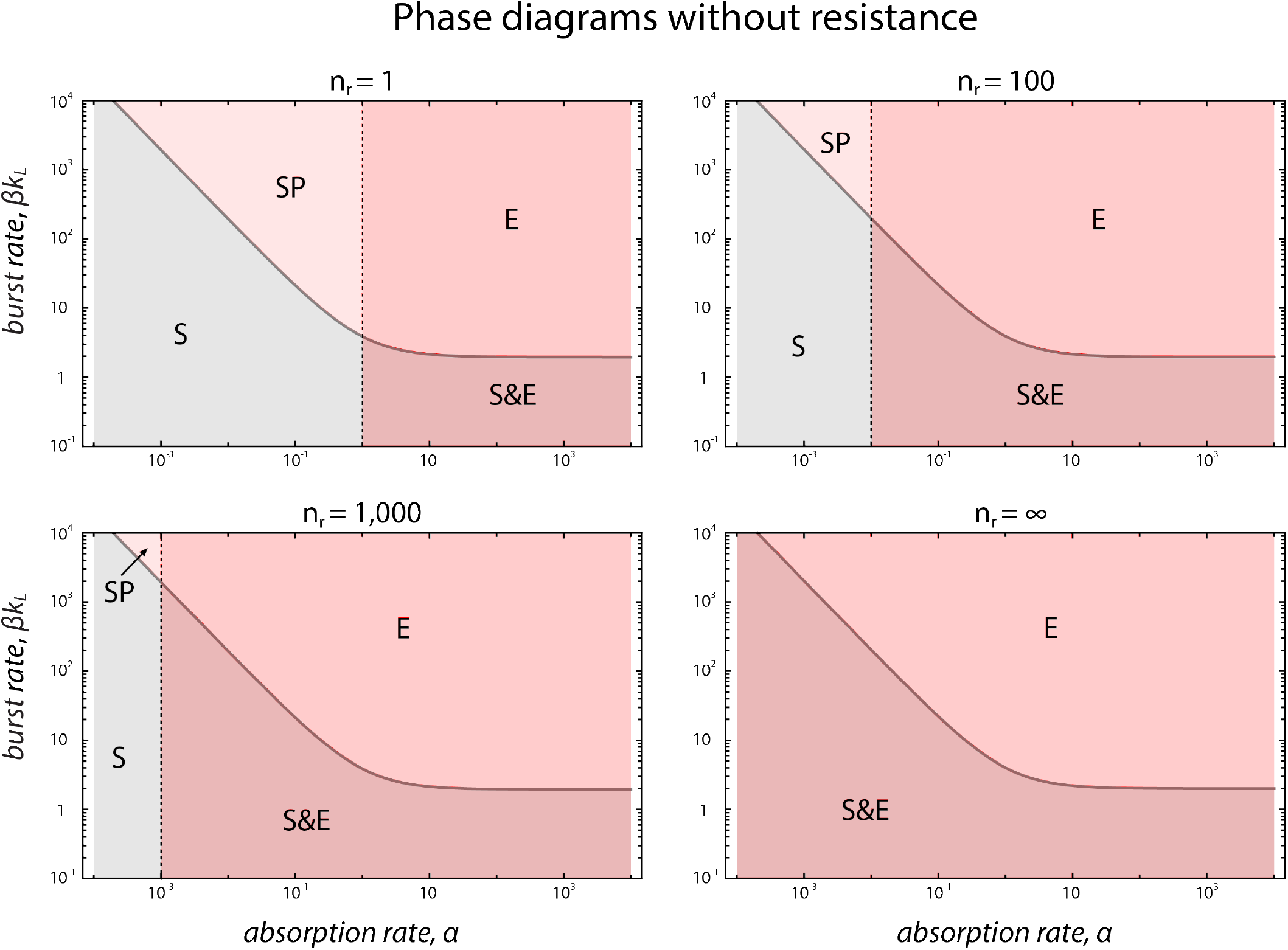
The phase diagrams without resistance, as a function of the number of receptors *n*_*r*_, phage burst rate *βk*_*L*_ and phage absorption rate *α*. The distinct stable outcomes correspond to the S, SP and E phases, as defined in the text. Vertical dotted line indicates *α* = *d*/*n*_*r*_. Host extinction is stable for *α* > *d*/*n*_*r*_. Parameters used are *d* = 1, *k*_*L*_ = 1 and *κ*′ = 0.

**FIG. S2.**
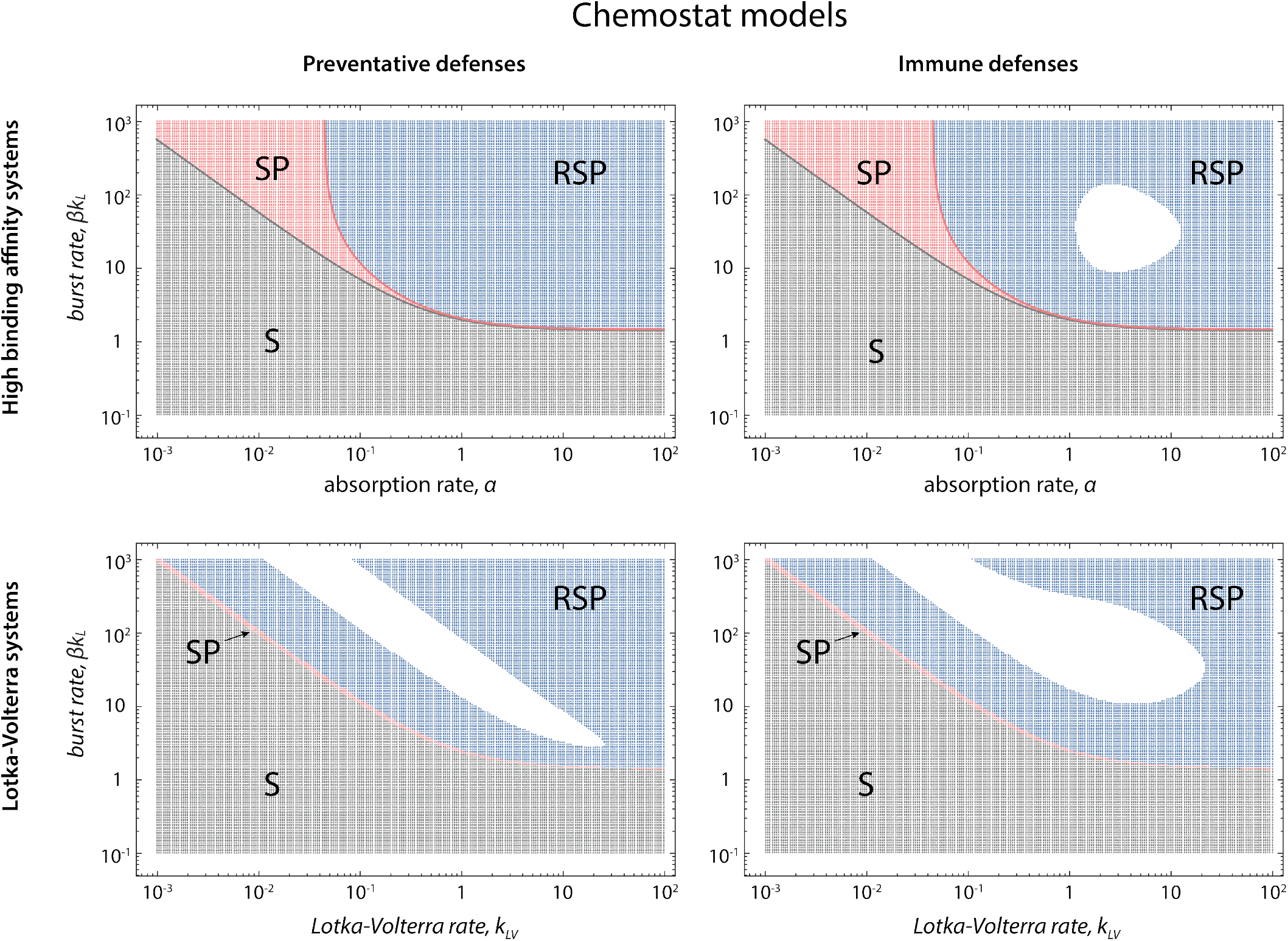
Phases of stable fixed points in chemostat models, numerically obtained. Left column - preventative defenses, right column - immune defenses, top row - infection rate of the form *k*_*I*_ = *αP*/(*A* + *P*), bottom row - Lotka-Volterra model with *k*_*I*_ = *k*_*LV*_ *P*. White regions indicate the absence of stable fixed points. Parameters used: *n*_*r*_ = 1, *d* = 1, *b* = 0.9, *k*_*L*_ = 1, *D* = 0.4, *r*_*in*_ = 1, *s* = 10^−4^.

**FIG. S3.**
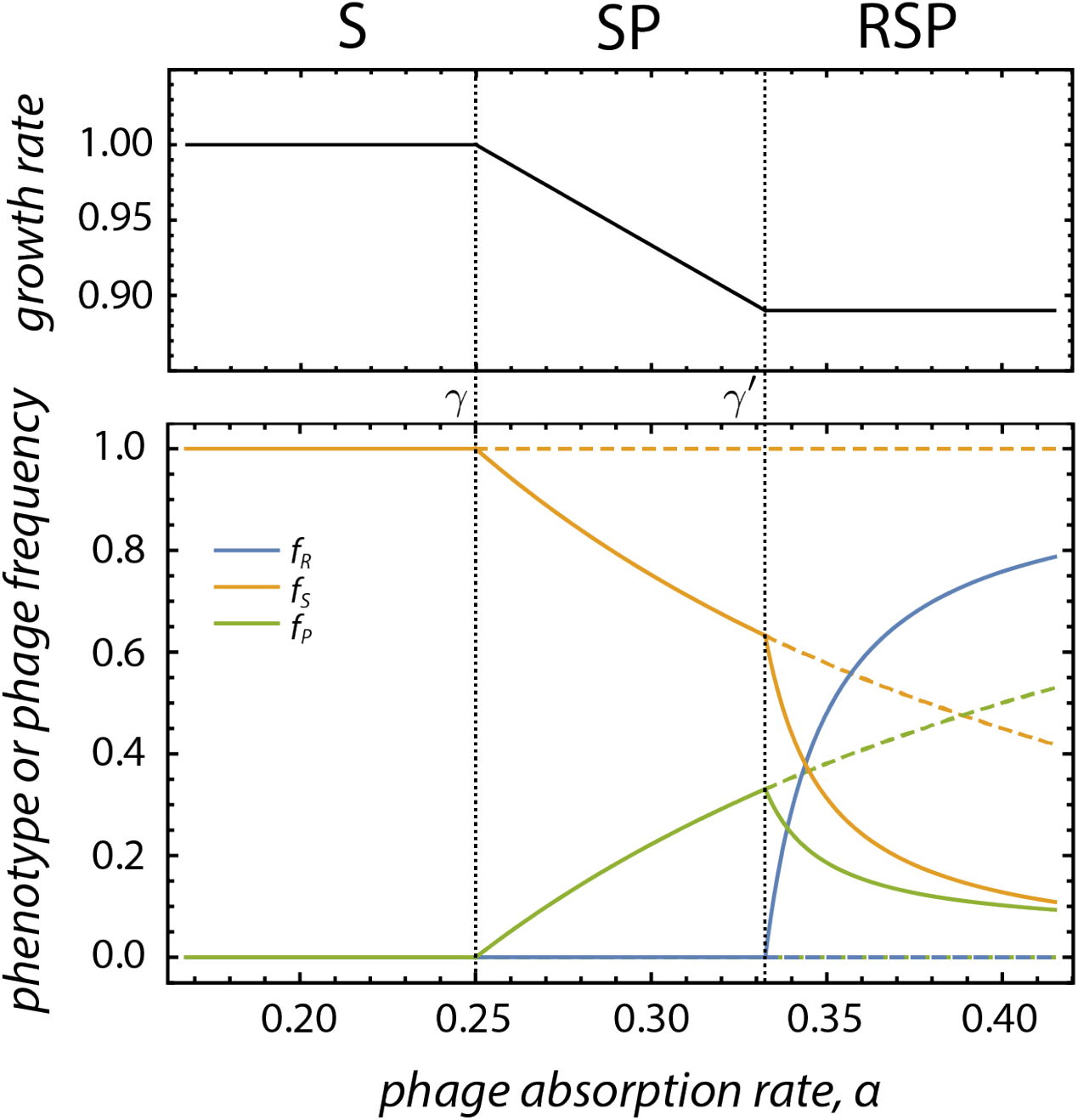
Plots showing the growth rate and phenotype and phage frequencies in the model of preventative defenses as *α* is varied across *γ* and *γ*′ at burst size *β* = 10. The system goes through a series of two transcritical bifurcations. Solid curves - stable fixed points, dashed curves - unstable fixed points. Remaining parameters are *n*_*r*_ = 1, *d* = 1, *b* = 0.9, *k*_*L*_ = 1, *s* = 0.01, *κ*′ = 0.

**FIG. S4.**
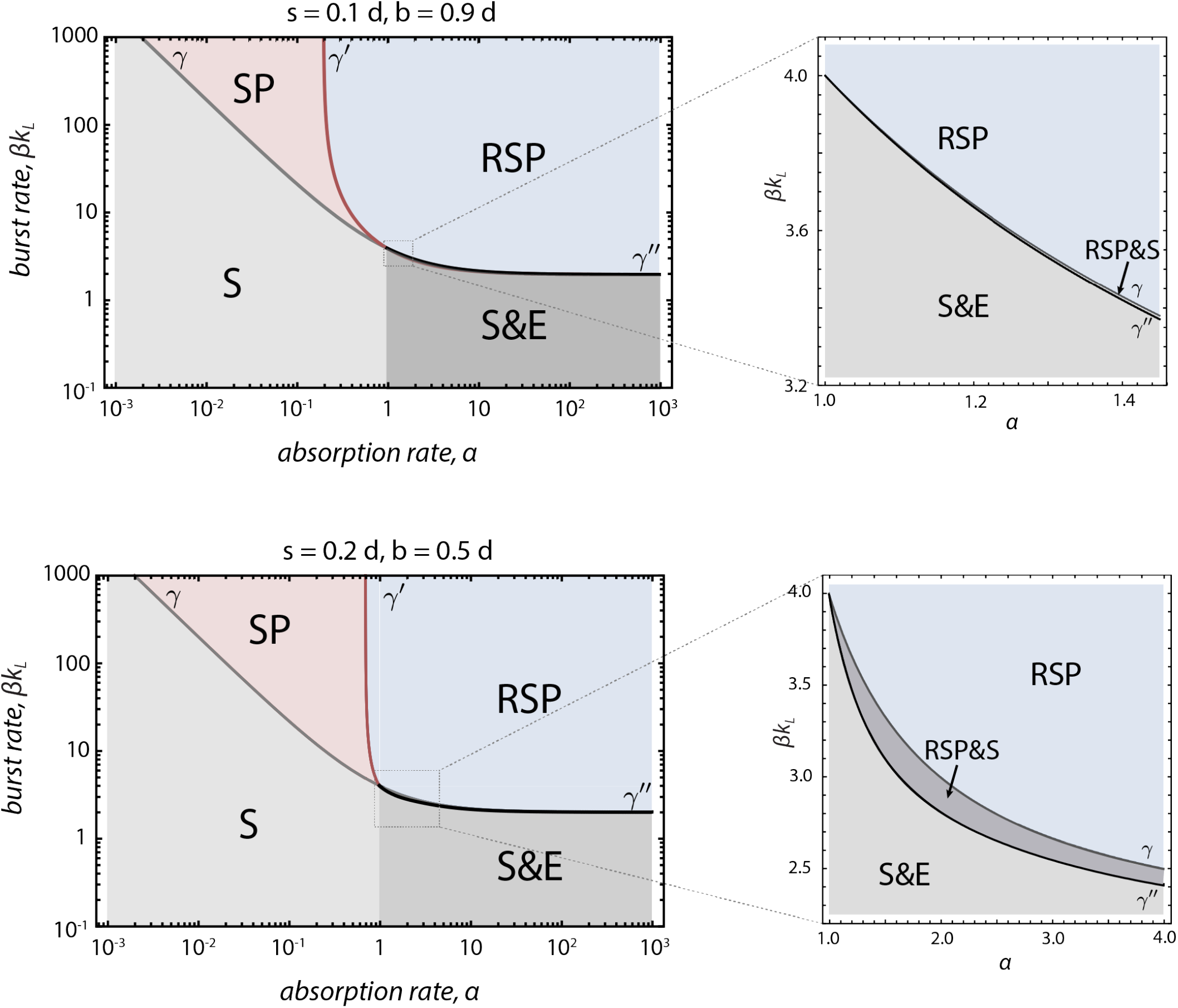
Phase diagrams for preventative defenses for larger switching rates and costs of defense mechanism, *d* − *b*. Insets show the separation of *γ* and *γ*″ for *α* > *d*. The RSP&S bistable region located between *γ* and *γ*″ curves becomes larger with the increase in switching rate and cost *d* − *b*. Remaining parameters are *n*_*r*_ = 1, *d* = 1, *k*_*L*_ = 1, *κ*′ = 0.

**FIG. S5.**
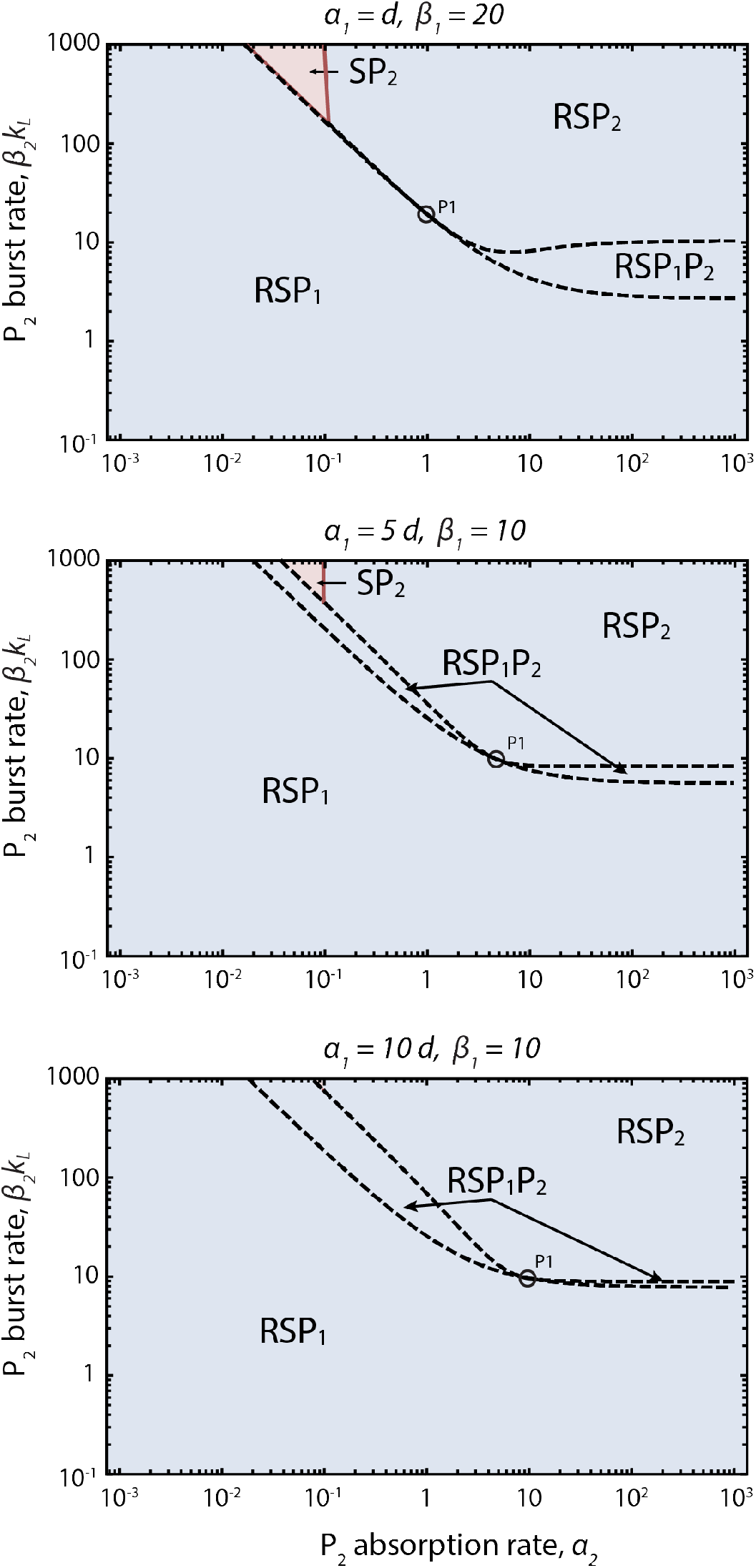
Phase diagram for two phage types, *P*_1_ and *P*_2_, shown for *P*_2_ parameters in the model of preventative defenses. *P*_1_ parameters (*α*_1_, *β*_1_) are located in the RSP phase. Dashed curves separate regions of single- and multi-phage phases. The pink region corresponds to the case where *P*_2_ drives *P*_1_ and the resistant phenotype extinct. The resistant phenotype switches at a rate *s* = 10^−3^. Size of the coexistence regions weakly depends on *s*. Remaining parameters are *d* = 1, *b* = 0.9, *k*_*L*_ = 1, *κ*′ = 0.

**FIG. S6.**
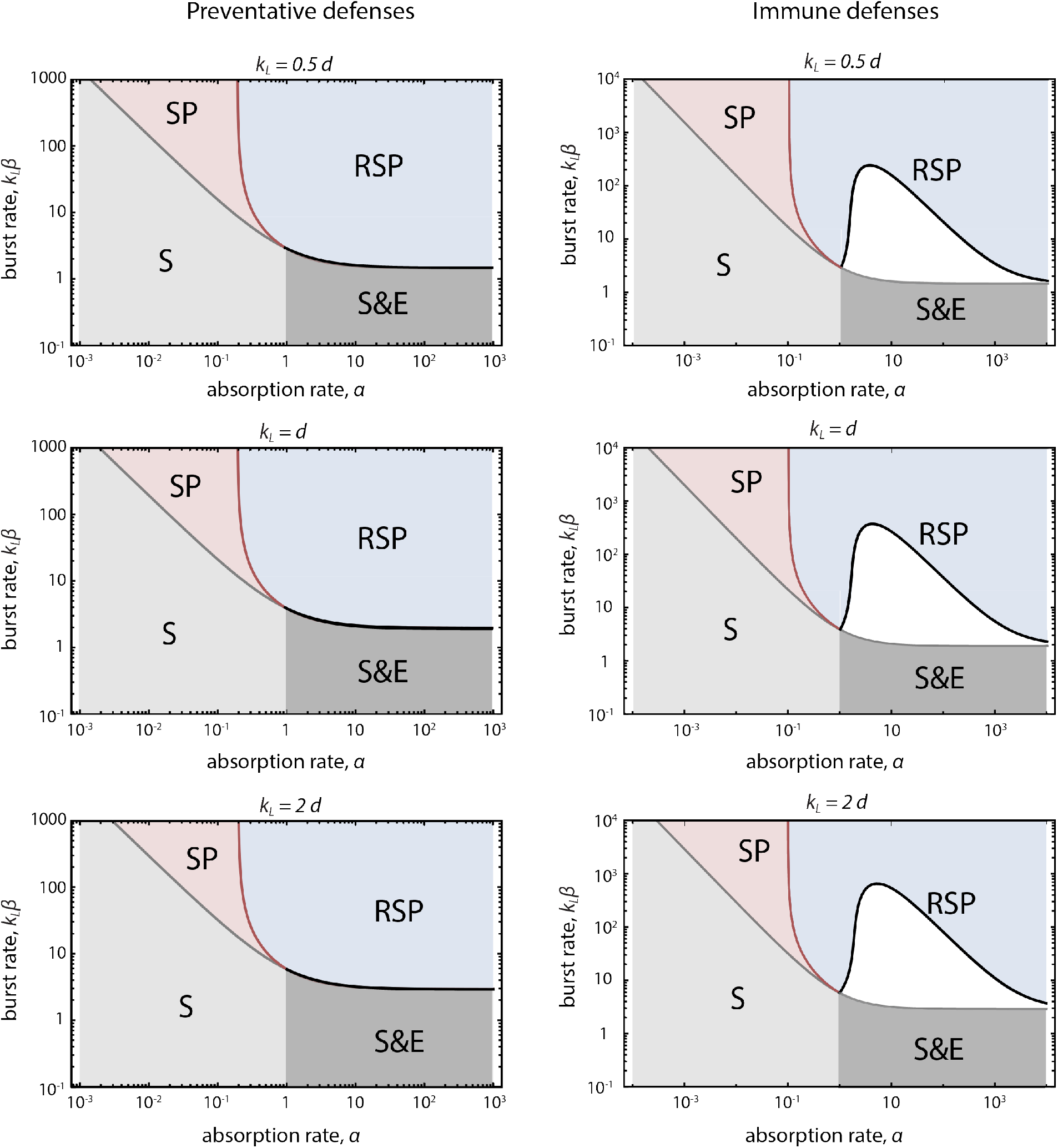
Phase diagrams for preventative and immune defenses, for different values of *k*_*L*_. Remaining parameters are *n*_*r*_ = 1, *d* = 1, *b* = 0.9, *s* = 10^−4^, *κ*′ = 0.

**FIG. S7.**
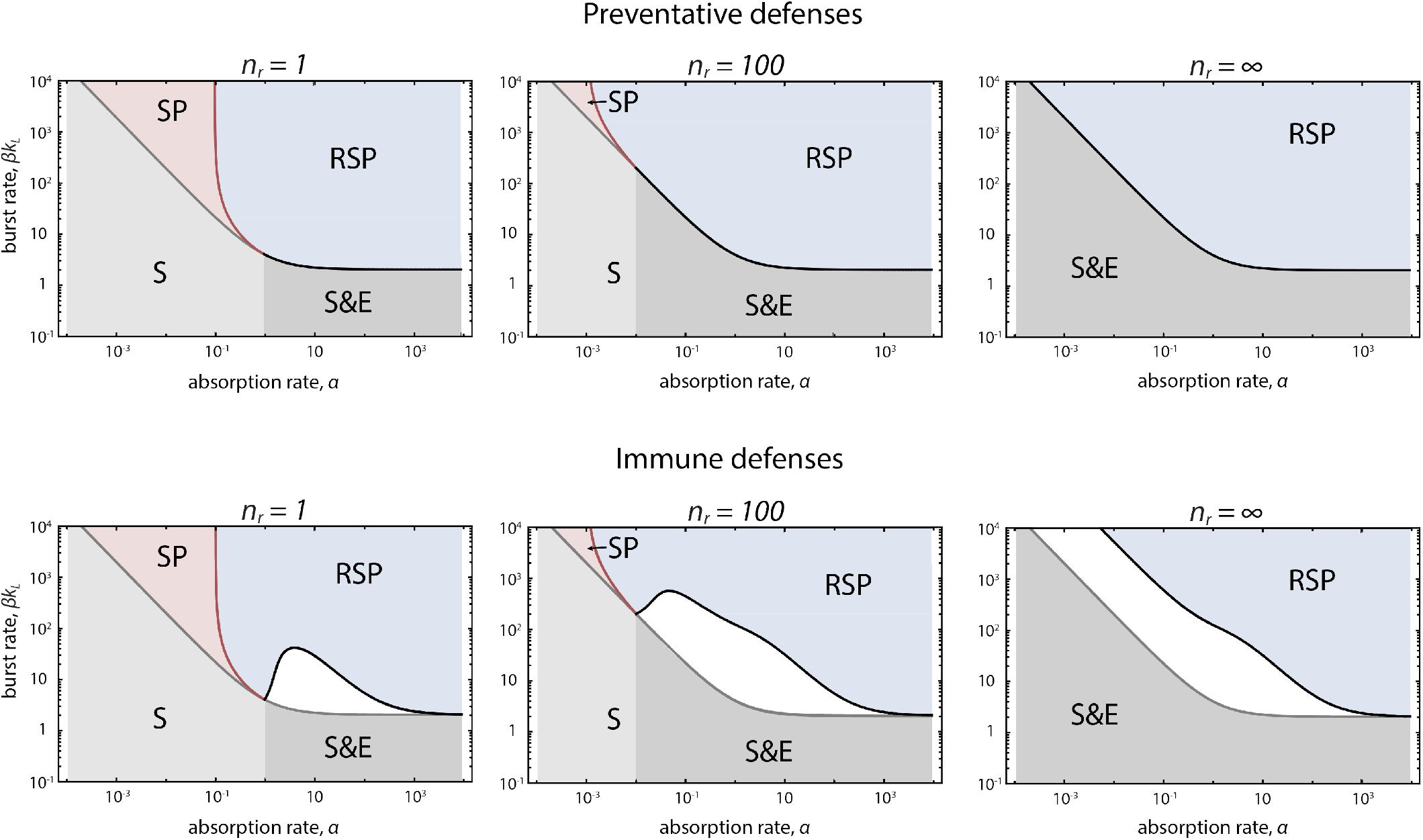
Phase diagrams for preventative and immune defenses for different values of *n*_*r*_. Remaining parameters are *d* = 1, *b* = 0.9, *s* = 10^−3^, *κ*′ = 0.

**FIG. S8.**
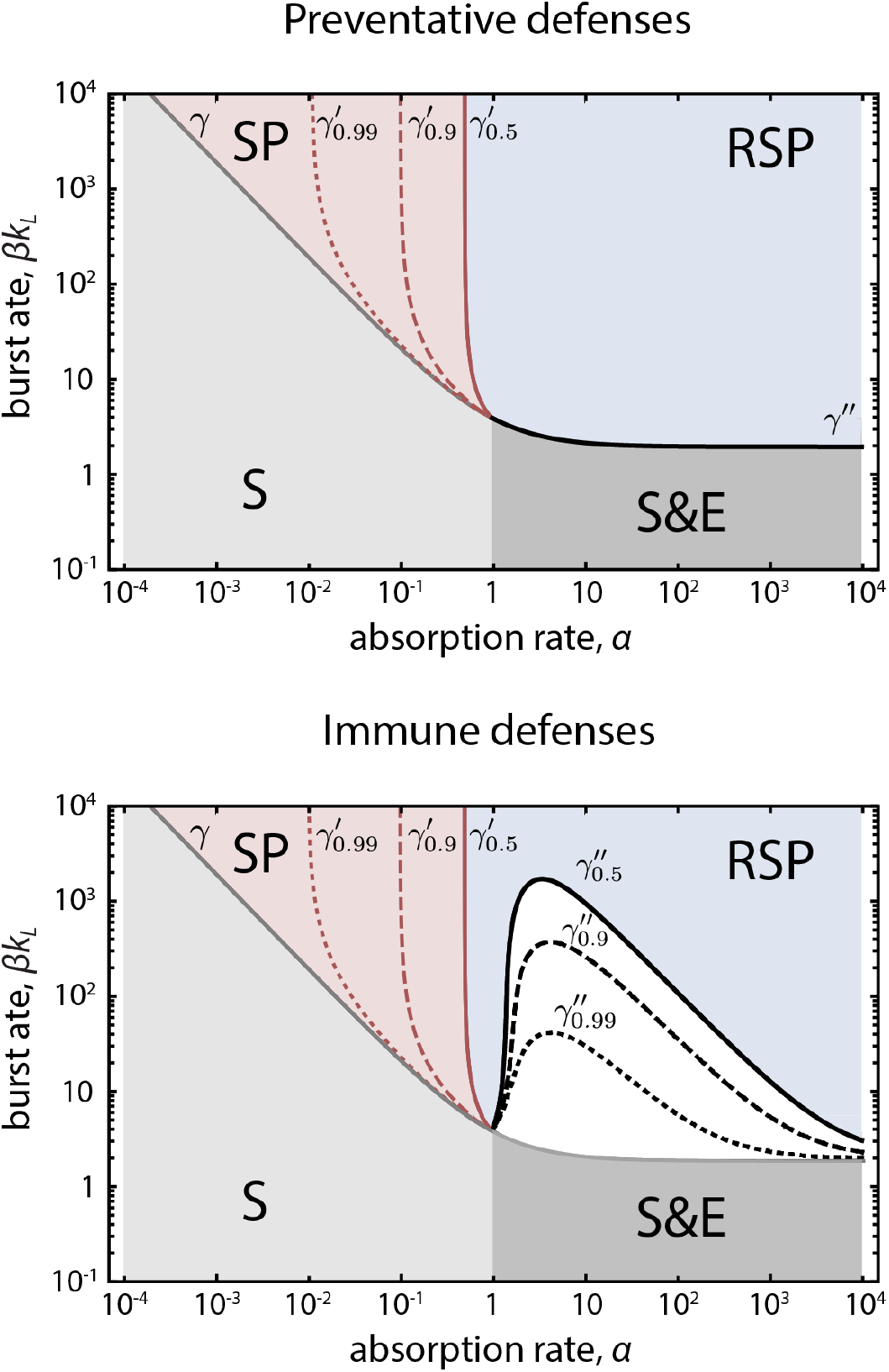
Phase diagrams for preventative and immune defenses, for different values of *b* (*b* = 0.5 - solid, *b* = 0.9 - dashed, *b* = 0.99 - dotted), corresponding to different fitness costs of defense. Bifurcation curves *γ*′ and *γ*″ are labelled according to their respective *b* values. For preventative defenses changes in *γ*″ occur within the width of the curve and feature a narrow RSP&S bistable region at the location of *γ*″ curve. The location of *γ* does not depend on *b*. Remaining parameters are *n*_*r*_ = 1, *d* = 1, *s* = 10^−4^, *κ*′ = 0.

**FIG. S9.**
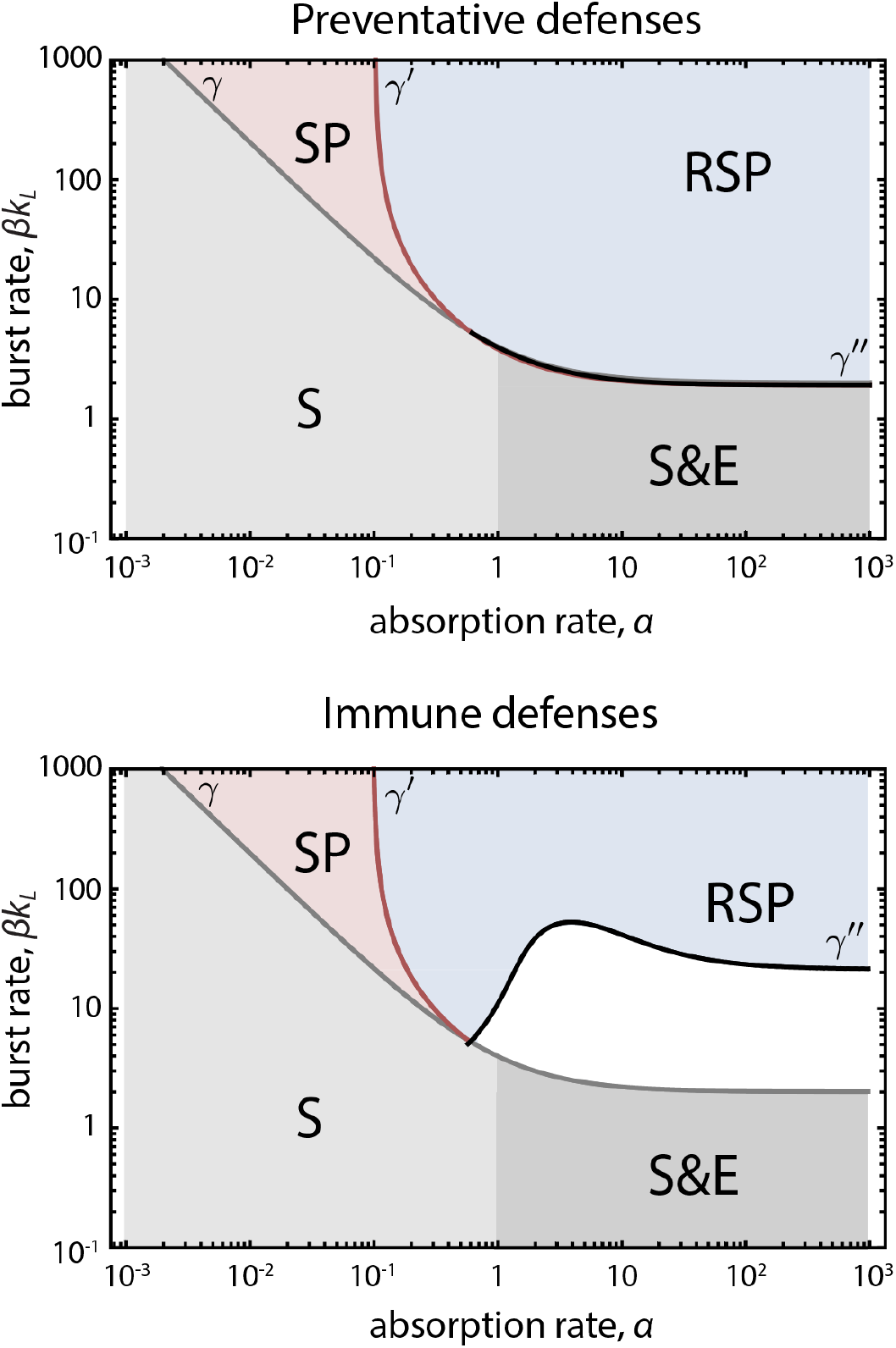
Phase diagrams for preventative and immune defenses where the infected phenotype does not absorb phage. Remaining parameters are *n*_*r*_ = 1, *d* = 1, *b* = 0.9, *k*_*L*_ = 1, *s* = 10^−3^, *κ*′ = 0.

**FIG. S10.**
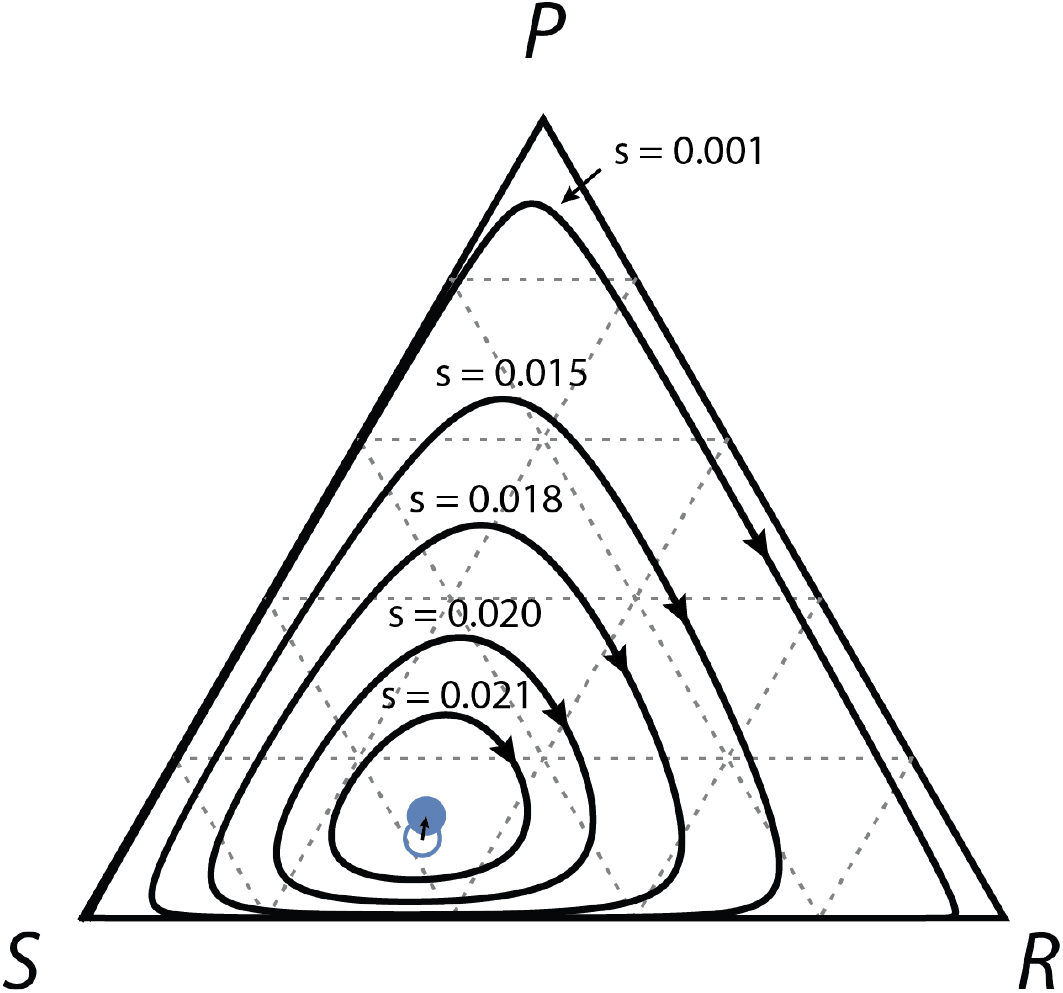
Ternary plot of limit cycle periodic orbits in the model of immune defenses for *α* = 1.5, *β* = 5 and *s* varied from 0.001 to 0.021. The unstable interior fixed point (empty blue circle) drifts with the increase of *s* in the direction of the arrow, and becomes stable for *s* ≈ 0.0216, which is represented with the filled circle. The periodic trajectories contract into the interior of the simplex and around the fixed point. Remaining parameters are *n*_*r*_ = 1, *d* = 1, *b* = 0.9, *k*_*L*_ = 1, *κ*′ = 0.

**FIG. S11.**
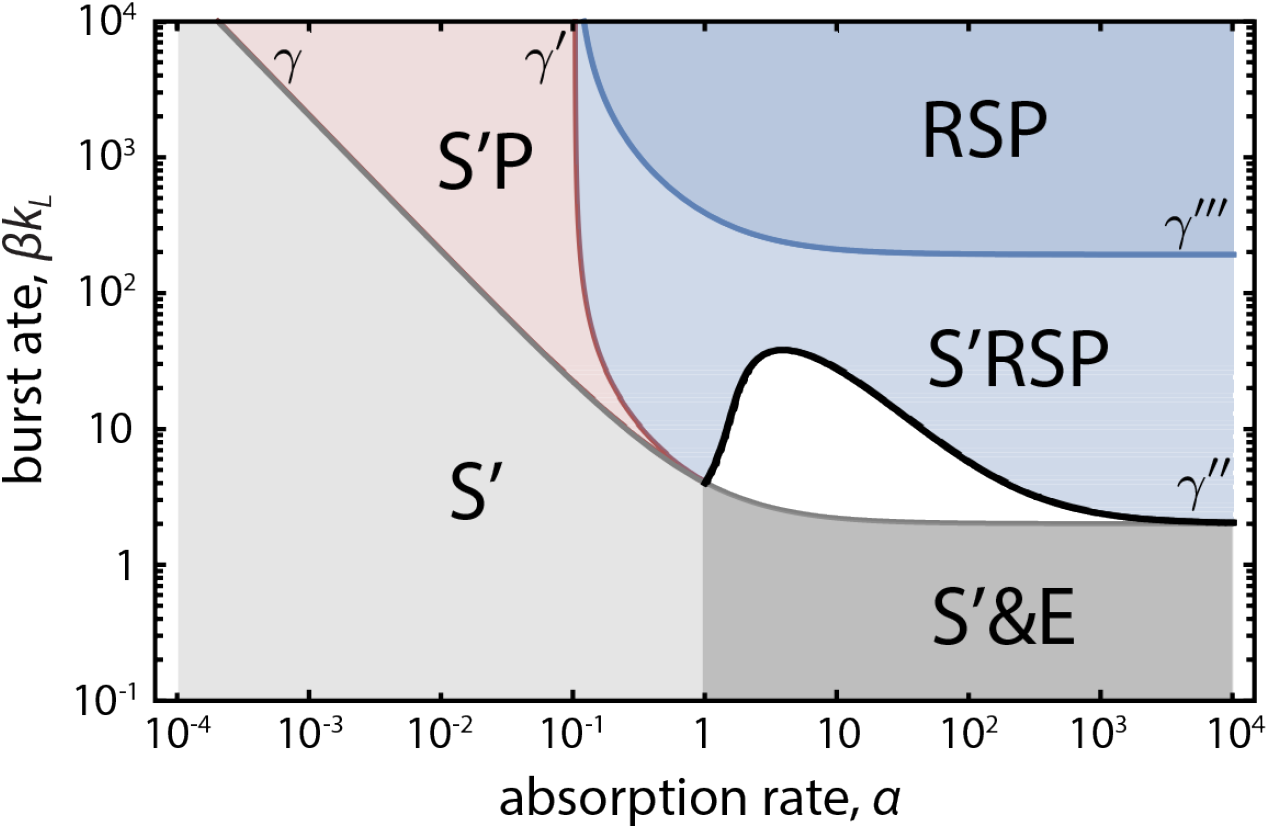
Phase diagram of the CRISPR spacer loss model. The resistant phenotype *R* generates the loss-of-spacer phenotype *S* which bears the cost of resistance without the benefit of immunity. *S*′ corresponds to a phage-sensitive strain that does not pay the cost of resistance. The resistant phase contains two regions, S’RSP and RSP, separated by *γ*‴ across which the system undergoes a transcritical bifurcation that eliminates *S*′ from the population. Coexistence of *S*′ with *RSP* indicates that *S*′ is effectively cheating off of the CRISPR system expressed by *R*. All the rates are in units of 1/division, with *s* = 10^−3^ and remaining parameters *n*_*r*_ = 1, *d*′ = 1, *b* = 0.9, *k*_*L*_ = 1, *κ*′ = 0.

